# Reporter system architecture affects measurements of noncanonical amino acid incorporation efficiency and fidelity

**DOI:** 10.1101/737197

**Authors:** K.A. Potts, J.T. Stieglitz, M. Lei, J.A. Van Deventer

## Abstract

The ability to genetically encode noncanonical amino acids (ncAAs) within proteins supports a growing number of applications ranging from fundamental biological studies to enhancing the properties of biological therapeutics. Currently, our quantitative understanding of ncAA incorporation systems is confounded by the diverse set of characterization and analysis approaches used to quantify ncAA incorporation events. While several effective reporter systems support such measurements, it is not clear how quantitative results from different reporters relate to one another, or which details influence measurements most strongly. Here, we evaluate the quantitative performance of single-fluorescent protein reporters, dual-fluorescent protein reporters, and cell surface displayed protein reporters of ncAA insertion in response to the TAG (amber) codon in yeast. While different reporters support varying levels of apparent readthough efficiencies, flow cytometry-based evaluations with dual reporters yielded measurements exhibiting consistent quantitative trends and precision across all evaluated conditions. Further investigations of dual-fluorescent protein reporter architecture revealed that quantitative outputs are influenced by stop codon location and N-and C-terminal fluorescent protein identity. Both dual-fluorescent protein reporters and a “drop-in” version of yeast display support quantification of ncAA incorporation in several single-gene knockout strains, revealing strains that enhance ncAA incorporation efficiency without compromising fidelity. Our studies reveal critical details regarding reporter system performance in yeast and how to effectively deploy such reporters. These findings have substantial implications for how to engineer ncAA incorporation systems—and protein translation apparatuses—to better accommodate alternative genetic codes for expanding the chemical diversity of biosynthesized proteins.

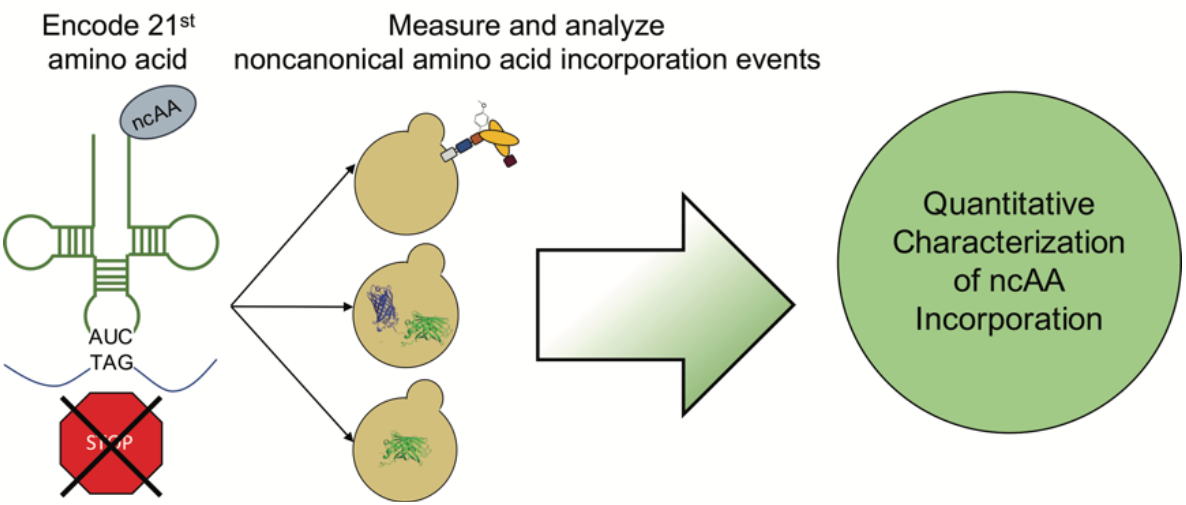

**Design, System, Application Paragraph:** On earth, the genetic code provides nearly invariant instructions for generating the proteins present in all organisms using 20 primary amino acid building blocks. Scientists and engineers have long recognized the potential power of altering the genetic code to introduce amino acids that enhance the chemical versatility of proteins. Proteins containing such “noncanonical amino acids” (ncAAs) can be used to elucidate basic biological phenomena, discover new therapeutics, or engineer new materials. However, tools for measuring ncAA incorporation during protein translation (reporters) exhibit highly variable properties, severely limiting our ability to engineer improved ncAA incorporation systems. In this work, we sought to understand what properties of these reporters affect measurements of ncAA incorporation events. Using a series of ncAA incorporation systems in yeast, we evaluated reporter architecture, measurement techniques, and alternative data analysis methods. We identified key factors contributing to quantification of ncAA incorporation in all of these categories and demonstrated the immediate utility of our approach in identifying genomic knockouts that enhance ncAA incorporation efficiency. Our findings have important implications for how to evolve cells to better accommodate alternative genetic codes.

## 1 Introduction

Genetically encoding noncanonical amino acids (ncAAs; also referred to as unnatural amino acids (uAAs), nonstandard amino acids (nsAAs), or non-natural amino acids (nAAs)) in proteins enables control over protein structure and function with atomic-level precision.^1–5^ Effective exploitation of ncAAs enhances our understanding of basic biology^6–8^ and provides opportunities for engineering new classes of materials^9^ and biological therapeutics.^2, 10–12^ Many of these applications require high efficiency, high fidelity ncAA incorporation and subsequent careful evaluation of such events. The use of mass spectrometry-based characterizations offers the highest level of rigor,^9, 13^ but lacks the throughput needed for initial screening and characterization. Additionally, mass spectrometry methods are not suitable for direct monitoring of incorporation events during protein translation. Another method for evaluating ncAA incorporation utilizes protein reporter systems in cells and cell-free translation systems as tools for understanding ncAA incorporation events, although the deployment of these systems varies widely. Even basic fluorescent reporters, where fluorescence is observed if a noncognate codon is suppressed, can possess drastic architectural differences between studies. These include the fluorescent protein variant utilized, the position within the reporter at which the ncAA is encoded, and data collection, analysis, and reporting methods.^14, 15^ Due in part to these structural variations, it remains unclear how differences in reporters affect quantitative characterizations of ncAA incorporation events. Using the protein translation apparatus to insert a ncAA into a protein sequence via codon suppression (stop codon,^16, 17^ 4-base codon,^18^ or codon containing unnatural bases^19^) is complex, and usually inefficient compared to wild-type protein translation. Numerous studies have shown that these inefficiencies can result from, but are not limited to, the activities of engineered aminoacyl-tRNA synthetase/tRNA pairs (i.e. orthogonal translation systems; OTSs),^9, 20, 21^ intracellular expression levels of OTS components,^22, 23^ activities of the ribosome,^24^ activities of elongation and release factors,^25, 26^ and the codon composition of the host genome.^27^ Elucidation of how these factors interact with one another is highly desirable for engineering translation apparatuses to accommodate alternative genetic codes. Integration of these observations and comparisons across studies requires a full understanding of how reporter systems and data analysis practices affect quantitative measurements of ncAA incorporation events.

Several reporter strategies for evaluating ncAA incorporation efficiency and fidelity have been described in the literature.^28, 29^ By far the most common approach is single-fluorescent protein reporters.^14, 15^ The primary advantage of these reporters is the easy-to-read fluorescent output, which in most cases is strongly correlated to the level of ncAA incorporation. However, readouts from these systems can be confounded by variability in intracellular plasmid levels or other processes that change reporter expression levels without altering suppression efficiency.^29^ In addition, it remains unclear how variations in the properties of these reporters, such as changing stop codon position, affects the quantitative evaluation of ncAA incorporation events. Recently described dual-fluorescent protein reporters, which consist of two fluorescent proteins with distinct spectral properties connected by a linker, have some inherent advantages over single-fluorescent protein reporters.^29^ Because these constructs provide a means of detecting both the expression level of the reporter (N-terminal fluorescent protein prior to codon for suppression) and full-length protein (C-terminal protein), variations in reporter system expression can be accounted for during analysis. Barrick and coworkers introduced the metrics “Relative Readthrough Efficiency” and “Maximum Misincorporation Frequency” for quantifying the efficiency and fidelity of ncAA incorporation, respectively, while normalizing for changes in reporter expression levels.^29^ Both single-and dual-fluorescent reporters support moderate to high throughput measurements with microplate readers and flow cytometry. One potential weakness of single- and dual-fluorescent reporters is that the folding time of fluorescent proteins may confound accurate determination of codon suppression efficiency. Recent reports have demonstrated the use of cell surface display systems for evaluating codon suppression events.^28, 30^ We recently showed that detection of the N- and C-termini of yeast-displayed constructs facilitates the use of the rigorous relative readthrough efficiency and maximum misincorporation frequency metrics described above. In addition, the surface accessibility of the reporter enables the use of chemical modifications to confirm the presence of a ncAA containing a specified reactivity, reminiscent of earlier residue-specific ncAA incorporation engineering work.^31^ Söll and coworkers implemented the use of an *E. coli* display system to screen for aminoacyl-tRNA synthetase (aaRS) variants that support ncAA incorporation based on full-length protein expression and selective chemical modification to identify the presence of a specific ncAA,^32^ but did not report quantitative measures of incorporation with this system. The use of epitope tags or conjugation reactions eliminate the potential for fluorescent protein folding rates to confound analysis. A primary drawback is the need to label the displayed constructs of interest with suitable detection reagents prior to quantitative evaluation.

Enzyme reporters of codon suppression that enable colorimetric readouts of enzyme activities have also been implemented in yeast and *E. coli*.^17, 33^ Like fluorescent reporters, these enzymes decouple codon suppression events from cell survival, and provide a means of evaluating relative levels of suppression activity. Coupling codon suppression events to cell survival has been utilized in both prokaryotic and eukaryotic cells.^17, 34^ These life-or-death assay formats support positive and negative selections and the ability to tune selection stringencies. On the other hand, these assays do not support quantitative measurements of ncAA incorporation efficiency or fidelity. The varied properties of the reporters described above suggests that each system has a role to play in discovering and evaluating ncAA incorporation systems. However, it remains unclear how results from distinct systems can be compared with one another due to significant differences in reporter system design, data collection, and data analysis. In this study, we investigated the performance of three types of reporter systems that support fluorescence-based measurements of ncAA incorporation efficiency and fidelity. Our work here is conducted in *S. cerevisiae*, the only organism in which quantitative measurements of ncAA incorporation efficiency and fidelity with single-fluorescent protein reporters,^15, 35, 36^ dual-fluorescent protein reporters,^28^ and display-based reporters^28^ have all previously been performed (Fig. 1). We compared reporters constructed in these three formats using flow cytometry and microplate-based measurements (when possible) to evaluate ncAA incorporation efficiency and fidelity. In our hands, flow cytometry-based measurements led to more precise measurements than microplate-based measurements across all systems tested. Examination of a series of ncAA incorporation events known to exhibit a range of efficiencies and fidelities yielded similar trends in each reporter format. However, observed levels of ncAA incorporation efficiency and fidelity varied as a both a function of the system used and the method of downstream analysis. Based on these results, we constructed a series of dual-fluorescent protein reporters to better understand the effects of varying the fluorescent proteins utilized, orientation of proteins within the reporter, and TAG codon location and number. We then investigated the utility of several reporters for assessing ncAA incorporation events in a series of yeast knockout strains harboring genomic deletions of nonessential genes known to affect protein translation. Multiple strains supported enhanced ncAA incorporation efficiency without apparent loss of fidelity. We also found that controlling for changes in wild-type reporter expression levels is critical to determining whether a genomic modification is attributable to changes in codon suppression. These findings highlight the utility of these reporters in evaluating how the protein translation apparatus can be engineered to better support the use of alternative genetic codes and should support genome engineering efforts to construct and evolve organisms that utilize such codes. Taken as a whole, our results provide important insights into how to effectively deploy reporter systems in search of ncAA incorporation systems that expand the chemical versatility of proteins.

**Fig. 1.**
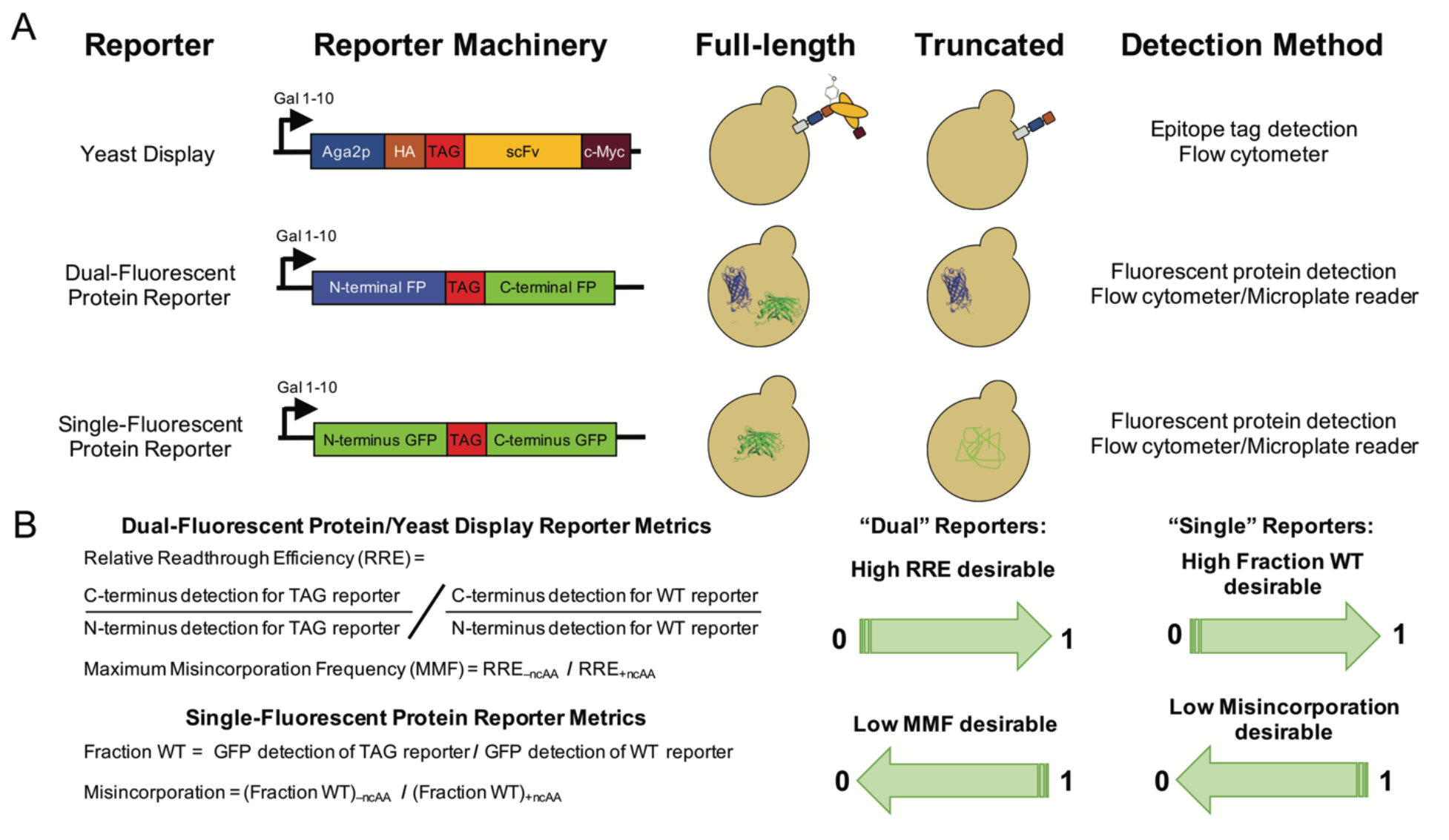
Reporter architectures, detection methods, and analysis methods. (A) Architectures of the three major types of reporters used in this work, expected behaviors in yeast, and detection methods for quantifying ncAA incorporation events. (B) Metrics used to determine ncAA incorporation efficiency and fidelity based on N- and C-terminal detection in “dual” reporters (yeast display, dual-fluorescent protein reporter) and full-length protein detection in “single” reporters (single-fluorescent protein reporter).

## 2 Materials and Methods

### 2.1 Materials

All restriction enzymes used for molecular biology were from New England Biolabs (NEB). Synthetic oligonucleotides for cloning and sequencing were purchased from Eurofins Genomics or GENEWIZ. All sequencing in this work was performed by Eurofins Genomics (Louisville, KY) or Quintara Biosciences (Cambridge, MA). Epoch Life Science GenCatch^TM^ Plasmid DNA Mini-Prep Kits were used for plasmid DNA purification from *E. coli*. Yeast chemical competent cells and subsequent transformations were prepared using Zymo Research Frozen-EZ Yeast Transformation II kits. *O*-methyl-L-tyrosine and *p*-azido-L-phenylalanine were purchased from Chem-Impex International, Inc. (catalog numbers 06251 and 06162, respectively).

### 2.2 Media preparation and yeast strain construction

The preparation of liquid and solid media was performed as described previously.^28^ The strain RJY100 was constructed using standard homologous recombination approaches and has been described in detail previously.^37^ The strain BY4741 (YSC1048) was purchased from Dharmacon. The BY4741 knockout strains from the Yeast Knockout Collection were obtained from the laboratory of Stephen P. Fuchs at Tufts University. The BY4705 strains (shown in Fig. S12) were obtained from the laboratory of Catherine Freudenreich at Tufts University (stock numbers 483 and 646 in the Freudenreich Lab) and were originally purchased from ATCC (*Saccharomyces cerevisiae* ATCC 200869^TM^).

### 2.3 Reporter plasmid construction

The pCTCON2-FAPB2.3.6, pCTCON2-FAPB2.3.6L1TAG, pCTCON2-RYG, and pCTCON2-RXG reporter constructs have been previously described.^28^ The pCTCON2-BXG reporter was cloned by replacing the RFP segment in pCTCON2-RXG with a BFP gene amplified from pBAD-mTagBFP2, obtained from the laboratory of Nikhil U. Nair at Tufts University and originally purchased from Addgene (Addgene plasmid # 34632; http://n2t.net/addgene:34632; RRID:Addgene_34632), by digesting both pCTCON2-RXG and the PCR-amplified BFP gene with EcoRI-HF and BamHI-HF (NEB), then ligating with T4 DNA ligase (NEB) to insert BFP. The pCTCON2-BXG-2TAG and pCTCON2-BXG-altTAG constructs were cloned by Gibson assembly with EcoRI-HF-and PstI-HF-digested pCTCON2-BXG. Primers were designed to introduce the alternative TAG codon at the first serine residue in the linker sequence between the BFP and sfGFP genes and revert the TAG codon at the end of the linker back to the tyrosine residue in the wild-type linker sequence.^29^ Primers for the 2TAG reporter introduced the TAG codon at the first serine residue in the linker and maintained the second TAG codon at the original location in pCTCON2-BXG. The pCTCON2-GXB and pCTCON2-GYB reporter constructs were cloned by amplifying BFP and GFP from pCTCON2-BXG with primers designed to maintain the same linker sequence with and without the TAG codon positioned at the original location in the linker, then cloned into EcoRI-HF-and BglII-digested pCTCON2 via Gibson assembly. The first amino acid in sfGFP, which was an alanine in the pCTCON2-BXG/BYG constructs, was reverted back to methionine. The pCTCON2-GFP constructs were cloned by amplifying pCTCON2-BXG with primers to revert the first residue in GFP back to methionine, then cloned into EcoRI-HF-and BglII-digested pCTCON2 via Gibson assembly. For the pCTCON2-GFP-TAG construct, additional primers were used to introduce a stop codon in place of tyrosine at the 151st amino acid position of the construct. The PCR products corresponding to the two sfGFP fragments before and after the 151st amino acid were cloned into pCTCON2 via Gibson assembly. pCTCON2-Aga1p-FAPB2.3.6 and pCTCON2-Aga1p-FAPB2.3.6L1TAG were constructed in two steps. First, the Aga1p gene was amplified from YIP SHRPa-Aga1p and cloned via Gibson Assembly into pCTCON2 digested with restriction enzymes AgeI and KpnI. YIP sHRPa-Aga1P was a gift from Alice Ting (Addgene plasmid # 73151; http://n2t.net/addgene:73151; RRID:Addgene_73151). The resulting plasmid, pCTCON2-Aga1p, was sequence verified. The FAPB2.3.6 and FAPB2.3.6L1TAG genes were amplified from pCTCON2-FAPB2.3.6 and pCTCON2-FAPB2.3.6L1TAG, respectively, and then cloned via Gibson assembly into the pCTCON2-Aga1p vector digested with restriction enzymes BamHI-HF and NheI-HF. The resulting plasmids, pCTCON2-Aga1p-FAPB2.3.6 and pCTCON2-Aga1p-FAPB2.3.6L1TAG, were sequence verified. pRS416-Aga1p-FAPB2.3.6 was constructed by amplifying the Aga1p-FAPB2.3.6 segment from pCTCON2-Aga1p-FAPB2.3.6 and using a Gibson Assembly to insert the fragment into pRS416 digested with restriction enzymes XbaI and SalI-HF. The sequence verification revealed a point mutation in the Aga1p gene that was subsequently removed. The TAG version of the pRS416-Aga1p-FAPB2.3.6 reporter was made by cloning in a TAG codon at the first position of the light chain of the scFv in the same position as the other yeast display reporter plasmids. Resulting plasmids were sequence verified. The promoter and BXG or BYG DNA fragments were amplified from pCTCON2-BXG and pCTCON2-BYG and cloned into XbaI-and SalI-HF-digested pRS416 via Gibson assembly for the pRS416-BXG and pRS416-BYG plasmids, respectively. pRS416-BXG-altTAG was constructed by amplifying the BFP-GFP DNA fragment with the alternative TAG codon from pCTCON2-BXG-altTAG and inserting it via Gibson assembly into pRS416 double digested with XbaI and SalI-HF. Resulting plasmids were sequence verified.

### 2.4 Suppressor plasmid construction

Suppressor plasmids pRS315-OmeRS^30, 37^ containing the tyrosyl OmeRS and pRS315-LeuOmeRS^28^ containing the leucyl OmeRS have been previously reported and characterized in detail.

### 2.5 Preparing noncanonical amino acid liquid stocks

All ncAA stocks were prepared at a 50 mM concentration of the L-isomer. DI water was added to the solid ncAA to approximately 90% of the final volume, and 6.0 N NaOH was used to fully dissolve the ncAA powder in the water by vortexing. Water was added to the final volume and the solution was sterile filtered through a 0.2 micron filter. OmeY solutions were pH adjusted to 7 using HCl. Filtered solutions were stored at 4°C for up to one week for AzF and two weeks for OmeY prior to use.

### 2.6 Yeast transformations, propagation, and induction

Reporter construct plasmids containing either a TRP1 (pCTCON2) or URA3 (pRS416) marker and aaRS/tRNA suppression plasmids pRS315-OmeRS and pRS315-LeuOmeRS (LEU2 marker) were transformed simultaneously into Zymo competent *S. cerevisiae* strains RJY100, BY4705 483, BY4705 646, BY4741, and the BY4741 deletion strains, plated on solid SD-SCAA media (either −TRP −LEU, −LEU −URA, or −TRP depending on the combination of plasmids), and grown at 30°C until colonies appeared (3 days).

Biological triplicates were used for all quantitative measurement experiments. All cells were grown and induced in tubes regardless of whether readthrough was assessed on a flow cytometer or on a plate reader. All liquid cultures were supplemented with a 100X penicillin/streptomycin to a final concentration of 1X (Corning 100X Penicillin:Streptomycin solution) to decrease the probability of contamination. To propagate samples with biological replicates and prepare them for induction, three separate colonies from each transformation were inoculated in 5 mL selective media and allowed to grow to saturation at 30°C (2−3 days). For cases where liquid colonies were already available, samples from saturated cultures stored at 4°C were pelleted and resuspended to an OD_600_ of 0.5−1.0 in 5 mL fresh media and allowed to grow to saturation overnight. Following saturation, the cultures were diluted to an OD_600_ of 1 in fresh media and grown at 30°C until reaching mid log phase (OD 2−5; 4−8 h). Cells were pelleted (5 min at 2,400 rpm) and resuspended to an OD_600_ of 1 in induction media (cells containing reporter construct only: SG-SCAA (−TRP); cells containing both reporter constructs and suppression constructs: either SG-SCAA (−TRP −LEU), SG-SCAA (−LEU −URA), or SG-SCAA (−TRP −LEU −URA), depending on the combination of suppressor and reporter plasmids in each yeast strain). To enable site-specific incorporation of ncAAs, induction media was supplemented with 1 mM final concentration of the L-isomer of the following ncAAs: *O*-methyl-L-tyrosine (pH 7) and *p*-azido-L-phenylalanine, and then induced at 20°C for 16 h.

### 2.7 Flow cytometry data collection and analysis

Freshly induced samples were labeled in 1.7 mL microcentrifuge tubes or 96-well V-bottom plates. Flow cytometry was performed either on an Attune NxT flow cytometer (Life Technologies) at the Tufts University Science and Technology Center or on a BD™ LSR II (BD Biosciences) at the Tufts University Flow Cytometry Core in the Jaharis Building. Labeling of induced yeast cultures with antibodies for detection of the N-and C-terminal epitope tags has been previously described in detail^28^ and was not modified for these experiments.

### 2.8 Plate reader data collection

To measure RFP and sfGFP levels for fluorescent protein reporters co-transformed with suppression constructs, 2 million cells per sample of freshly induced cells were pelleted (5 min at 2,400 rpm) and washed 3 times with 1x PBSA in 96-well V-bottom plates and then transferred to Corning 96-well clear bottom black-walled microplates for fluorescence measurements. Cultures containing pCTCON2-FAPB2.3.6 with no suppressor were used to measure the autofluorescence of the cells (cell blank), as they were not expected to exhibit any RFP, BFP, or sfGFP detection. All samples, including the cell blank, were run in biological triplicate and resuspended in 200 μL room temperature PBSA before measurements were taken. Fluorescence and OD measurements were performed using a SpectraMax i3X microplate reader (Molecular Devices, LLC., San Jose, California). OD readings were taken as end point measurements at 600 nm. RFP, GFP, and BFP readings were taken as end point measurements with RFP excitation and emission wavelengths set at 550 nm and 675 nm, respectively. The GFP excitation and emission wavelengths were set to 480 nm and 525 nm, respectively. The BFP excitation and emission wavelengths were set to 399 nm and 456 nm, respectively.

### 2.9 Calculating RRE and MMF

Detailed methods for flow cytometry RRE and MMF data analyses for yeast-displayed reporter constructs, including error propagation, has been described previously.^28^ Dual-fluorescent reporter RRE and MMF analyses were performed similarly, replacing HA and c-Myc detection with N-terminal and C-terminal fluorescent protein detection. Fraction of wild-type (Fraction WT) and misincorporation of the sfGFP reporter were calculated by replacing the dual detection with single detection of sfGFP and comparing the TAG-containing constructs to wild-type sfGFP expression under the same media conditions (i.e. in the presence or absence of ncAA) using the equations provided in Fig. 1B. Both this work and our previous report use the Microsoft Excel function “STDEV” to determine standard deviation of samples measured in biological triplicate.

Microplate reader data analysis was performed using Microsoft Excel. The fluorescence from each sample was normalized by the sample’s respective OD and then averaged across the biological triplicates. The average normalized fluorescence of the cell blank triplicates was then subtracted from the normalized fluorescence sample average to corrected for yeast cell autofluorescence. For dual fluorescent protein reporters, both the N-terminal and C-terminal proteins were taken into account for RRE and MMF calculations, whereas the fraction WT and misincorporation of the sfGFP reporter were determined using single detection and the equations from Fig. 1B.

### 2.10 RRE error as a percent of the magnitude

To determine error as a percent of the magnitude, the error-propagated standard deviations were divided by the magnitude of the relative readthrough efficiencies. These fractions were then converted to percentages and reported in Table S2 and S3.

### 2.11 Alternative analysis methods

Alternate analyses of flow cytometry data were performed using FlowJo and Microsoft Excel. For each sample collected on the flow cytometer, the overall population was gated for live, single cell events to exclude doublet and triplet data from the downstream analysis. From these single cell populations we performed the first set of alternate efficiency calculations (“Single Cell Population”) by averaging the median fluorescence intensity (MFI) data from the C-terminal fluorescent protein and taking the standard deviation of the biological triplicate (Fig. S2, S14, S21, S28). The second alternate analysis, “Reporter Expressing Cells + Background Subtraction,” utilized MFI values for C-terminal detection in cells expressing the reporter (i.e. cells demonstrating above-background levels of N-terminal fluorescent protein detection in the case of the dual reporters) and MFI values in cells not expressing a reporter (nonexpressing cells). We then subtracted the nonexpressing-cell MFI values from MFI values for the subset of cells with above-background levels of N-terminal fluorescent protein detection. For the single-fluorescent protein reporters, the population was gated into cells exhibiting above-background GFP fluorescence and background-level GFP fluorescence. The MFI of the backgroup-level GFP fluorescence population was subtracted from the above-background GFP population to obtain background-subtracted MFIs. The averages and standard deviations of the biological triplicates were then used to calculate propagated standard error (Fig. S3, S15, S22, S29). Equations used to propagate error are as previously described.^28^ The last alternate analysis (“Reporter Expressing Cells + BG Subtraction Normalized to WT Reporter”) takes the values as described in the second analysis method (“Reporter Expressing Cells + Background Subtraction”) and reports the TAG-containing constructs as a fraction of the respective wild-type construct (without a TAG codon and in the presence of no ncAAs) efficiency (Fig. S4, S16, S23, S30). All three of these alternate analyses were calculated using median fluorescence intensity as well as mean fluorescence intensity (Fig. S6−S8, S18−S20, S25−S27, S32−S34).

Microplate reader alternate analyses were performed using Microsoft Excel. For the first of the alternate microplate calculations (OD Normalized), the fluorescence intensity of the C-terminal fluorescent protein measured in each sample was normalized to the sample’s respective OD and then averaged with the associated biological triplicates. In the case of the single-fluorescent protein reporter, the fluorescence intensity of sfGFP was measured in place of the C-terminal fluorescence measurement (Fig. S9). For the second microplate reader analysis (OD Normalized + Background Subtraction), the normalized average fluorescence of the cell blank triplicates was subtracted them from the average normalized fluorescence of the sample to correct the fluorescence signals for yeast cell autofluorescence (Fig. S10). The last of the alternate microplate reader analyses, “OD Normalized + BG Subtraction Normalized to WT Reporter,” reported the previous TAG-containing construct efficiencies as a fraction of the related wild-type construct (without a TAG codon and in the presence of no ncAAs) efficiency (Fig. S11).

Finally, data from the dual reporters presented in Fig. 2 and Fig. 5 were subjected to another analysis (Reporter Expressing Cell Population) in which the median fluorescence intensity of the N-terminal signal (i.e. HA epitope detection or N-terminal fluorescent protein detection) was reported for the wild-type construct in the absence of any ncAA as well as for the TAG constructs in the presence of OmeY, AzF or in the absence of ncAA (Fig. S5, S17, S24, S31).

**Fig. 2.**
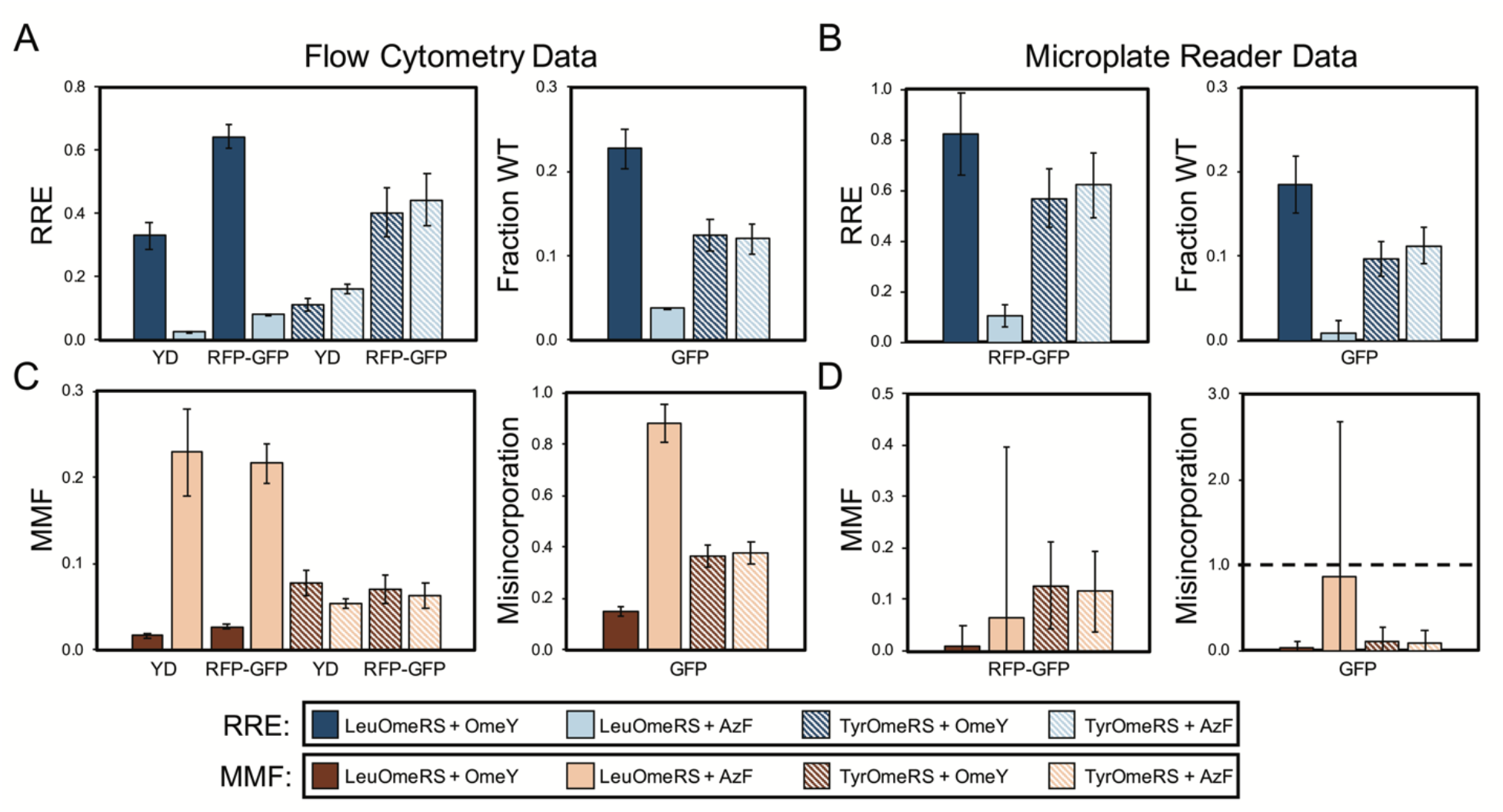
Quantification of ncAA incorporation efficiency and fidelity for four orthogonal translation system-ncAA combinations. (A) Flow cytometry data quantifying relative readthrough efficiency (RRE) for “dual” reporter systems (yeast display (YD), RFP-GFP dual-fluorescent protein reporter) and fraction of wild-type reporter (Fraction WT) for GFP single-fluorescent protein reporter. (B) Microplate reader data quantifying RRE for RFP-GFP reporter and Fraction WT for GFP reporter. (C) Flow cytometry data quantifying maximum misincorporation frequency (MMF) or misincorporation for RRE and Fraction WT measurements, respectively, reported in (A). (D) Microplate reader data quantifying MMF or misincorporation for RRE and Fraction WT measurements, respectively, reported in (B). All conditions were evaluated using end point measurements in biological triplicate. YD, yeast display.

## 3 Results and Discussion

### 3.1 Framework for evaluating the performance of reporters of noncanonical amino acid incorporation in yeast

We initiated our studies of the performance of ncAA reporter systems by implementing reporters with architectures previously described in the literature (Fig. 1A): a yeast display-based reporter, a dual-fluorescent protein reporter, and a single-fluorescent protein reporter. Both the yeast display-based reporter and dual RFP-GFP reporter are identical to the reporters that we have described previously,^28, 29^ with a TAG codon between two epitopes or proteins that support fluorescence detection. Moreover, the sfGFP used in the single-fluorescent protein reporter is genetically identical to the GFP in RFP-GFP. The amber codon in sfGFP was introduced at Y151, which is a commonly used permissive site for amber suppression.^14, 23^ All three of these reporters are under the control of the inducible Gal 1-10 promoter and allow for the modular introduction of aminoacyl-tRNA synthetase/tRNA pairs comprising the orthogonal translation system (OTS) machinery. These reporters were introduced into yeast along with each of two previously reported OTSs that were originally engineered to incorporate *O*-methyl-L-tyrosine in response to TAG codons in yeast: an *E. coli* tyrosyl-tRNA synthetase/tRNA^Tyr^_CUA_ pair (TyrOmeRS)^38^ and *E. coli* leucyl-tRNA synthetase/tRNA^Leu^_CUA_ pair (LeuOmeRS)^39^ where the LeuOmeRS variant was modified further to contain a T252A mutation.^28^ In previous work, we found that TyrOmeRS supports moderate levels of ncAA incorporation in the presence of either OmeY or *p*-azido-L-phenylalanine (AzF), with low but detectable levels of TAG codon readthrough in the absence of ncAAs. LeuOmeRS supports high levels of ncAA incorporation with OmeY, very low levels of ncAA incorporation with AzF, and essentially undetectable levels of readthrough in the absence of ncAAs. Thus, induction of the reporters under these different conditions provides a large expected range of readthrough efficiencies and fidelities for evaluation of reporter system performance.

All measurements of ncAA incorporation efficiency were conducted using end point measurements collected in biological triplicate. Following collection of data for all samples and controls using a flow cytometer (all reporters) or a microplate reader (fluorescent protein reporters), we determined ncAA incorporation efficiency and fidelity (Fig. 1B; see Materials and Methods for further details). For dual reporters (yeast display, dual-fluorescent protein reporters), we used the relative readthrough efficiency (RRE) metric and maximum misincorporation frequency (MMF) metric as previously introduced by Barrick and coworkers.^29^ These metrics are normalized to wild-type control constructs while also accounting for possible perturbations from the presence of the ncAA or changes in expression of the reporter during induction. For single-fluorescent protein reporters, which do not support the use of RRE and MMF, we determined the fraction of wild-type sfGFP (Fraction WT) expression (Fig. 1B). For all measurements, the data we collected enabled us to investigate the effects of using other analysis methods (for example, fraction of wild-type GFP expression in RFP-GFP). Taken together, these reporter architectures, OTSs, and analyses provide a rigorous framework for evaluating the properties of reporters and their effects on ncAA incorporation efficiency and fidelity.

### 3.2 Comparisons of conventional reporter architectures

Fig. 2 depicts side-by-side comparisons of yeast display reporter, dual-fluorescent protein reporter, and single-fluorescent protein reporter measurements of ncAA incorporation efficiency and fidelity using the framework described above. Qualitatively, data collected with all reporters show the expected trends for ncAA incorporation efficiency and fidelity: LeuOmeRS +AzF < TyrOmeRS +OmeY ≅ TyrOmeRS +AzF < LeuOmeRS +OmeY. In addition, all measurements determined via flow cytometry exhibit a higher level of precision (relative to standard error) in comparison with microplate reader-based measurements. Flow cytometry measurements provide errors that are between 2% and 19% of the magnitude of the RRE, whereas measurements on the microplate reader provided errors on the range of 19% to over 100% (Table S2). For flow cytometry-based measurements, a similar level of error was observed for reporters based on either yeast-displayed proteins or fluorescent proteins, indicating that with flow cytometry these distinct reporters can all be used to measure ncAA incorporation efficiency and fidelity with high precision.

Because RRE and MMF were only first used as metrics of ncAA incorporation processes in 2017, we conducted a series of alternative data analyses in order to explore the effects of data analysis methodologies on quantitative outputs. Consistent with strategies reported in the literature, we considered only the C-terminal reporter signal for each construct and used this as the basis for determining the “fraction of wild-type behavior.” For flow cytometry data, we determined the median and mean of the C-terminal reporter signals on gated populations consisting of all single cells, or on only the population of cells exhibiting evidence of reporter expression (see Materials and Methods for further details on analysis). Fig. (S2 – S11) depict the results of our alternative analyses of the data collected for Fig. 2. For all three reporter architectures, trends in efficiency data are generally consistent with those determined in Fig. 2 across the 4 OTS-ncAA combinations considered here. However, there are some cases in which calculated values of efficiency differ from the results depicted in Fig. 2. As an extreme example, the incorporation efficiency of the LeuOmeRS-OmeY combination determined by yeast display ranges from approximately 30 to 50 percent depending on the analysis method, greater than a 1.5-fold variation. However, in many other cases, these variations are within calculated error and thus insignificant.

Measurements made with single-fluorescent protein reporters make the implicit assumption that the expression levels of reporter constructs do not change significantly when evaluating different OTS-ncAA combinations. To evaluate this assumption, we plotted the values of the N-terminal signal levels detected in all samples characterized in Fig. 2 using the Reporter Expressing Cell Population approach. Under these conditions, reporter expression levels are reasonably consistent, but can exhibit greater than 20% variability between samples in the same data series. This level of variation seems to be tolerable, as all reporter architectures considered here are able to distinguish between the different OTS-ncAA combinations considered in this work. However, the RRE measurement framework eliminates the risk of inadvertently neglecting changes in reporter system expression levels. While variations are low under the conditions evaluated in this section, changes in expression can become quite significant when evaluating incorporation events in different strains (in some cases, greater than 2-fold variation in reporter expression levels; see Section 3.5), suggesting that some caution should be exercised when using single-fluorescent protein reporters.

A final noteworthy observation is that for a given OTS/ncAA induction condition, the calculated values of ncAA incorporation efficiency and fidelity depend on the specific reporter system used. Since ncAA incorporation efficiency and fidelity determined with all three reporters follow the same trends across all conditions examined, this indicates that the specific architecture of the reporter system dictates the quantitative values determined in a given experiment. This observation motivated the design of reporters containing subtle variations in architecture to better understand how these variations effect quantitative metrics of ncAA incorporation efficiency and fidelity.

### 3.3 Variation of fluorescent proteins used in dual-fluorescent protein reporters

Given the comparable precision of display-and fluorescence-based reporters, we evaluated several dual-fluorescent protein reporters in which only the identities of the fluorescent proteins utilized were changed to investigate the role of fluorescent protein folding properties on reporter performance. Previous work raises the possibility that the long half maturation time of RFP in the RFP-GFP system could confound accurate determination of RRE and MMF (RFP maturation half-time in solution is on the order of one hour).^40^ To evaluate this possibility, we replaced RFP with blue fluorescent protein (BFP)^41^ to create a BFP-GFP dual-fluorescent protein reporter analogous to the RFP-GFP reporter (Fig. 3A). BFP has previously been reported to exhibit a maturation half-time in solution of approximately 12 minutes.^41^ To our surprise, the performances of the two reporter systems were comparable to one another despite the different folding times of the N-terminal fluorescent proteins (Fig. 3B). For the BFP-GFP system, we conducted measurements of RRE and MMF on multiple flow cytometers and observed similar quantitative values of efficiency, fidelity and error (Fig. S1; only one flow cytometer with optics suitable for evaluating the RFP-GFP system was readily available). We also switched the order of the BFP-GFP system to place sfGFP, which has a reported folding half-time in solution of under 1 minute,^42^ in the N-terminal position while maintaining the structure of the linker (Fig. 3A). Flow cytometry readouts with the resulting GFP-BFP reporter indicate that RRE and MMF values are similar to the RFP-GFP and BFP-GFP systems, but the precision of the measurements appears to be lower. In the case of LeuOmeRS with OmeY we observed that the calculated error of GFP-BFP was 43% of the RRE magnitude, whereas the error in the RFP-GFP and BFP-GFP systems was only 5.6% and 15% of the magnitude, respectively (Fig. 3B; Table S3). This trend also persisted for the other OTS-ncAA combinations evaluated (Table S3). It is not immediately clear why relocating a protein with an extremely fast folding rate to the N-terminus of a dual-fluorescent reporter should be detrimental to the performance of the reporter. In any case, these observations suggest that the fluorescent proteins in the dual reporter format cannot be treated as completely modular entities.

**Fig. 3.**
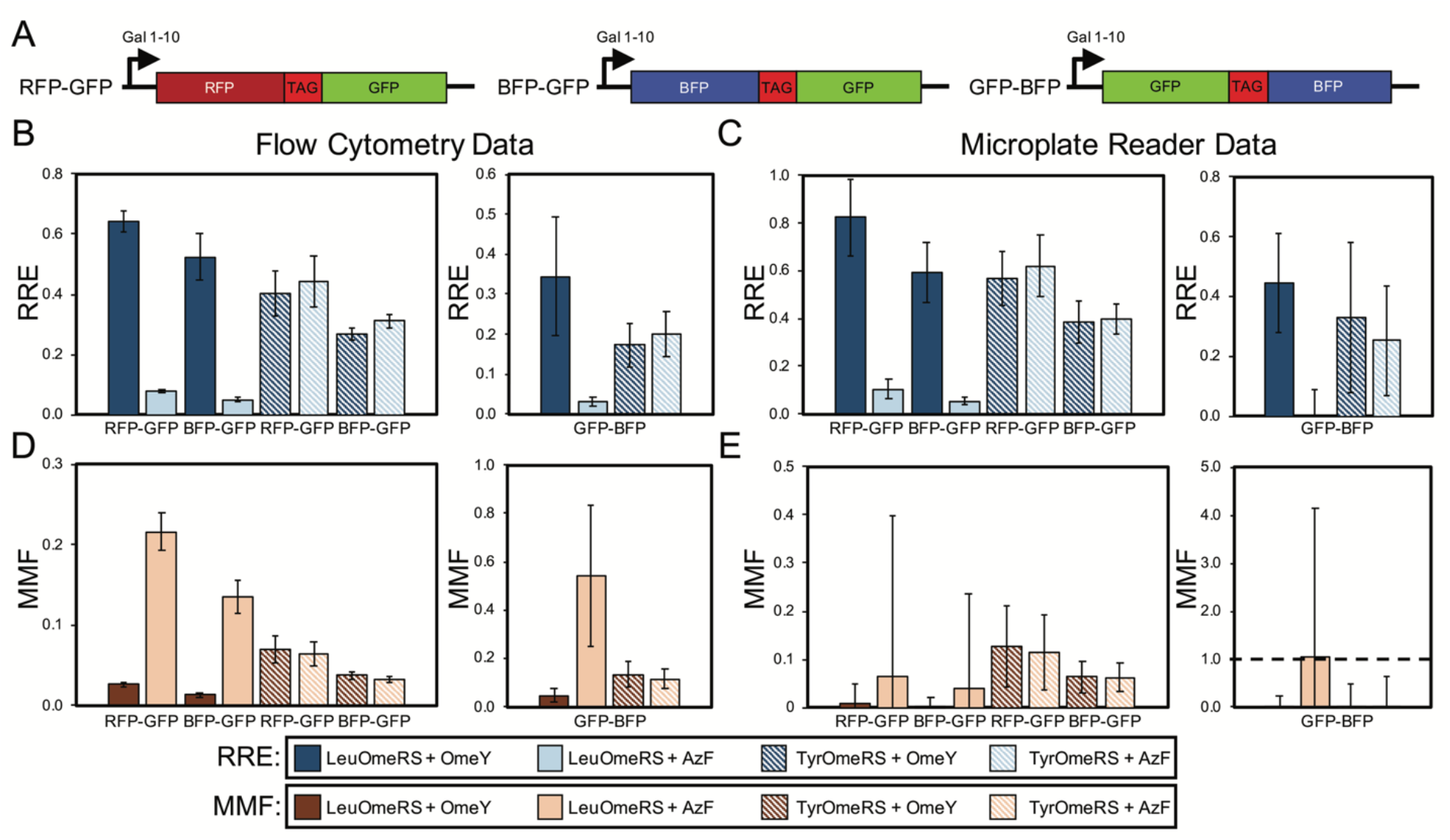
Evaluation of dual-fluorescent protein reporter performance as a function of fluorescent protein identity and orientation. (A) Architectures of RFP-GFP, BFP-GFP, and GFP-BFP reporters. The reporters contain identical linkers. (B) Relative readthrough efficiency determined via flow cytometry measurements with 4 OTS-ncAA combinations. Data for RFP-GFP and BFP-GFP was collected on an LSR-II flow cytometer, and data for GFP-BFP was collected with an Attune NxT flow cytometer (Fig. S1 indicates that data collected on these two instruments is equivalent). Note that the RFP-GFP data shown in Fig. 2 is being shown again to enable a direct comparison with the other dual-fluorescent protein constructs evaluated here. (C) RRE determined via microplate reader measurements for the same series of OTS-ncAA combinations as in (B). (D) Maximum misincorporation frequency determined via flow cytometry for RRE measurements reported in (B). (E) MMF determined via microplate reader for RRE measurements reported in (C). All conditions reported here were evaluated using end point measurements in biological triplicate.

Our flow cytometry measurements with all three dual-fluorescent reporters allow for reliable differentiation between the LeuOmeRS-ncAA and TyrOmeRS-ncAA incorporation events evaluated here. However, in several cases, the larger calculated error for experiments performed on the microplate reader makes reliable determination of differences in performance between OTS-ncAA combinations challenging (Fig. 3C; see Materials and Methods for calculation details). This observation is consistent with the data shown in Fig. 2, our own previous report,^28^ and that of Barrick and coworkers,^29^ where online measurements over the course of multiple hours with varying numbers of technological replicates per condition were used to reduce error during determination of RRE and MMF. For the remainder of our studies, we report only flow cytometry data due to its support of high-precision determination of ncAA incorporation efficiency and fidelity. Since the RFP-GFP and BFP-GFP dual reporters exhibit similar performance for the four OTS-ncAA combinations tested here, we chose to use BFP-GFP reporter variants to investigate the effects of altering stop codon positioning in the following section.

### 3.4 Variation of stop codon position and number in BFPRGFP dual-fluorescent protein reporter

Studies of nonsense suppression events in cells from several organisms have revealed that the context of a TAG codon, that is the bases flanking the codon, can affect stop codon readthrough efficiency.^43–47^ These observations and more direct studies of the role of the bases flanking TAG codons targeted for ncAA incorporation in *E. coli*^48^ highlight the need to investigate this issue further. Here, we examined a series of BFP-GFP reporters containing TAG codons at different positions for several reasons. First, for the same OTS-ncAA combination, the BFP-GFP reporter appears to support higher levels of ncAA incorporation in comparison to our yeast display reporter (Fig. 2). Second, in a previous study, we used the yeast display reporter to investigate three permissive stop codon locations within our antibody-based reporter and found ncAA incorporation efficiency to be consistent across the three positions, but lower than the BFP-GFP reporter efficiency observed in this study.^28^ Fig. 4A depicts variants of the BFP-GFP reporter in which the position of the stop codon has been moved to a different position within the linker (Alt-TAG) and in which two stop codons have been introduced into the vector simultaneously (2-TAG). We selected the first serine residue (TCC) to minimize the number of bases to be altered to facilitate TAG codon introduction and to avoid consecutive or near-consecutive TAG codons in the 2-TAG case. The RRE and MMF results in Fig. 4B clearly demonstrate that moving the position of the TAG codon to the alternate position (Alt-TAG) results in drastically lower suppression efficiencies with all combinations of OTSs and ncAAs tested. Introduction of two stop codons into the linker (at “standard” and “alternative” positions) further lowers measured ncAA incorporation efficiencies, although these efficiencies remain within a factor of two of the Alt-TAG efficiency values. Taken together with our previous observations,^28^ these data suggest that, at least in yeast, the original TAG codon position in the BFP-GFP (and RFP-GFP) reporter is especially permissive with respect to ncAA incorporation. The drastic differences between ncAA incorporation efficiencies observed here highlight the importance of characterizing and understanding reporter systems, as seemingly small changes in stop codon positioning can have a large impact on reporter output. A high level of “permissiveness” in a reporter may also obscure the detection of minute enhancements in ncAA incorporation efficiencies caused by genetic modification. To gauge the validity of this conjecture further, we conducted studies using strains from a single-gene knockout collection and various reporter systems to determine which reporters would support the detection of altered ncAA incorporation efficiency.

**Fig. 4.**
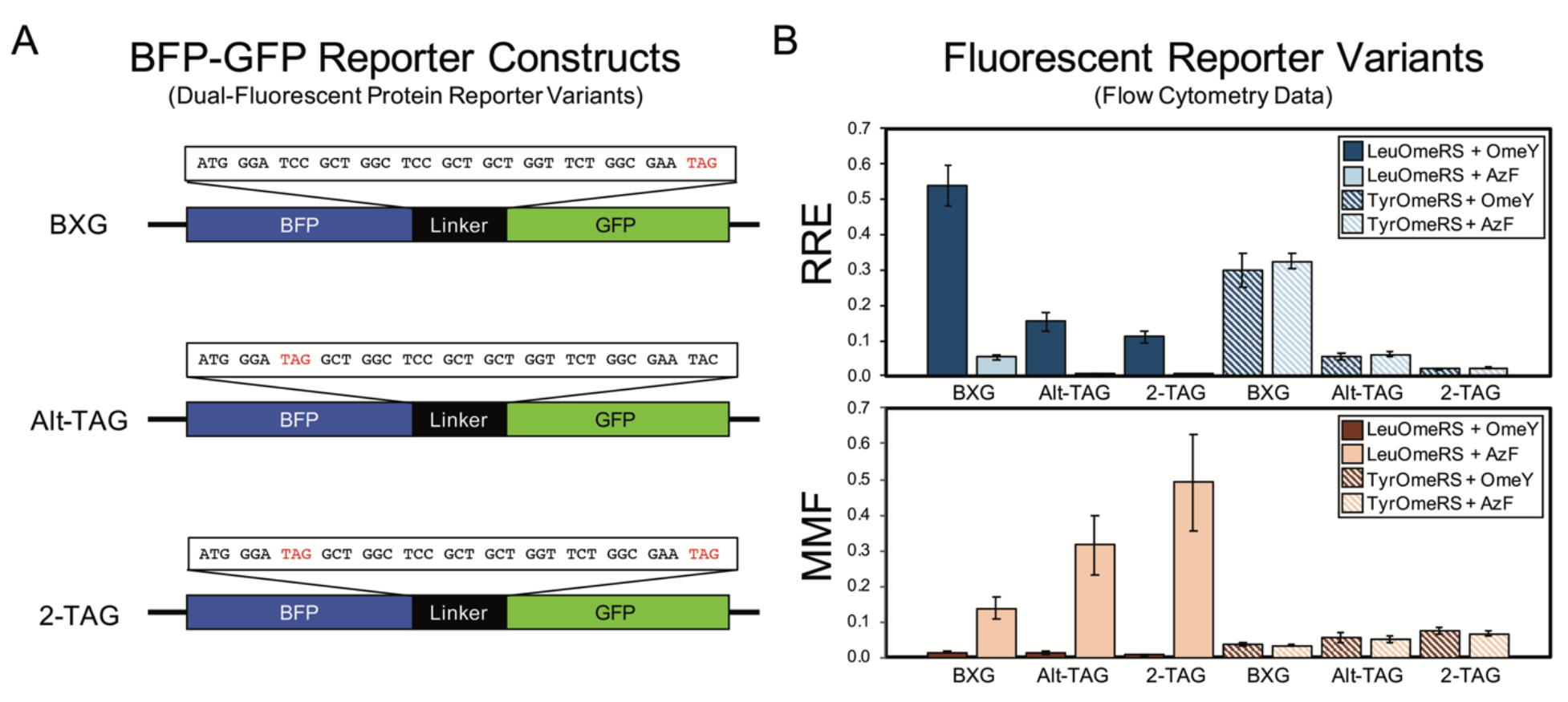
Evaluation of dual-fluorescent protein reporter performance as a function of stop codon position within linker. (A) Structures of BFP-GFP linker variants. (B) Relative readthrough efficiency and maximum misincorporation frequency determined via flow cytometry measurements with a series of four OTS-ncAA combinations. All conditions reported here were evaluated using end point measurements in biological triplicate.

### 3.5 Characterization of single-gene knockouts using reporter systems

Genomic modifications ranging from single-gene knockouts^26^ to complete overhaul of the genome^27^ can enhance ncAA incorporation via codon suppression events. Therefore, we wanted to determine whether the reporters characterized in this work would be suitable for evaluating ncAA incorporation efficiency and fidelity in yeast strains possessing several genotypes. We selected six strains from the haploid yeast knockout collection (YKO) containing a deletion in a nonessential gene known to be associated with efficiency of termination of protein translation.^49^ These strains and the parent BY4741 strain were co-transformed with OTSs and reporter systems and evaluated for ncAA incorporation efficiency and fidelity (Fig. 5). Because the strains of the knockout collection contain deletions of LEU2 and URA3, but not TRP1, we moved the BXG and Alt-TAG reporter systems from a pCTCON2 plasmid backbone into a pRS416 (URA3 marker) backbone. In addition, we prepared a “drop-in” version of our yeast display reporter within the pRS416 backbone suitable for use in any yeast strain (in contrast to conventional Aga1p-Aga2p yeast display, where Aga1p is integrated into the genome while Aga2p is encoded on pCTCON2. Fig. S12 provides a comparison of drop-in versus conventional yeast display using pCTCON2-based reporter plasmids, and indicates that the two display formats yield similar measurements of ncAA incorporation events (the yeast display strain RJY100 contains a genomically integrated URA3 marker, preventing use of pRS416-based plasmids in this strain). In BY4741, we observed that the LeuOmeRS-OmeY OTS-ncAA combination, which consistently yielded the highest relative readthrough efficiency in the pCTCON2 backbone (Fig. 2 – 4), exhibits similar RRE values to the TyrOmeRS-OmeY and TyrOmeRS-AzF OTS-ncAA combinations in the pRS416 backbone for all three reporters (Fig. 5A, 5B, Fig. S13 – S34). While clearly reproducible, the reason for this apparent shift in performance is not clear. This unexpected change in relative performance of OTS-ncAA combinations upon switching plasmid backbones further emphasizes the significant changes in reporter system performance that can result from seemingly inconsequential alterations to the reporters.

**Fig. 5.**
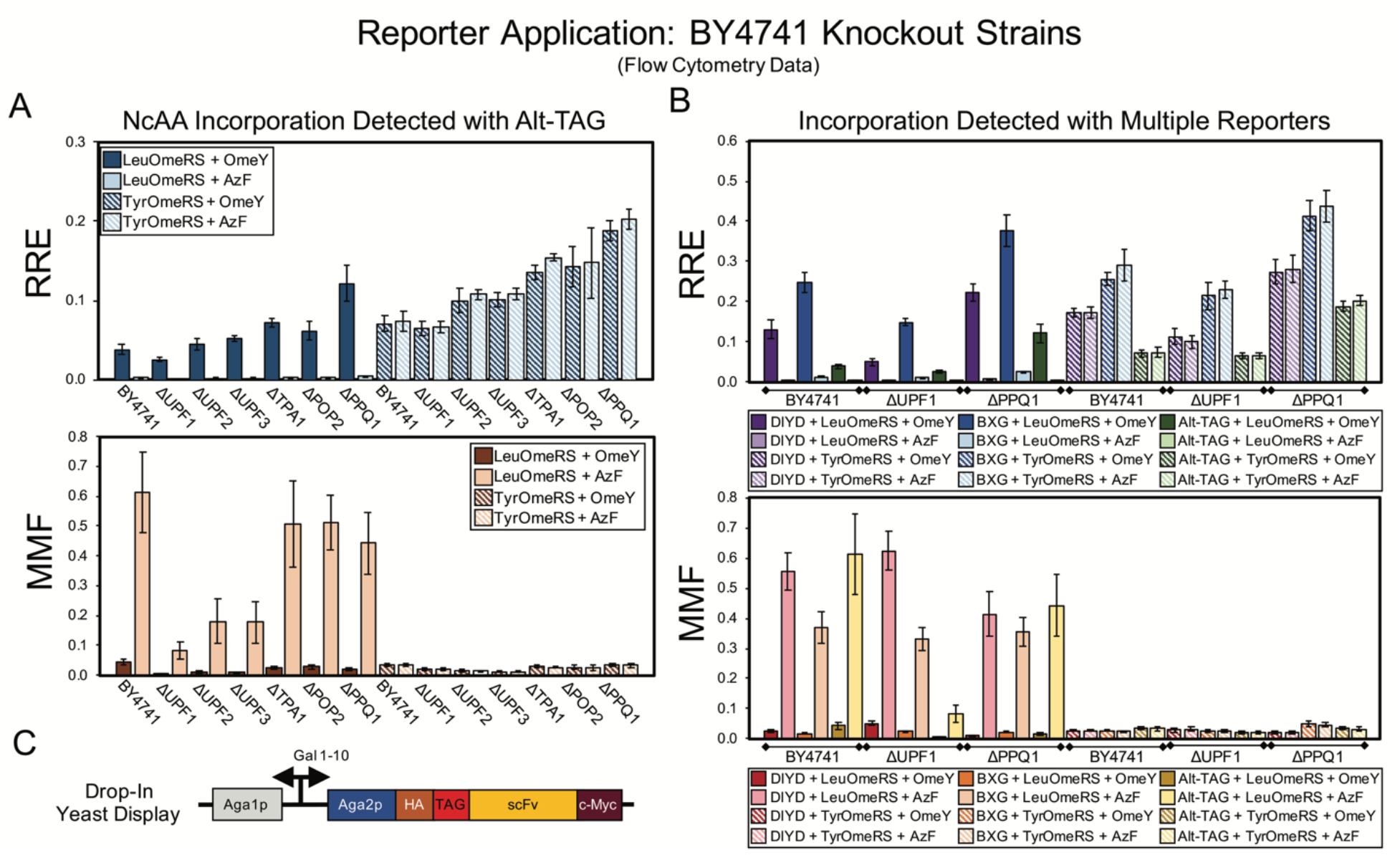
Evaluation of ncAA incorporation events in a series of single-gene knockout strains using two dual-fluorescent protein reporters and a drop-in yeast display (DIYD) reporter. (A) Measurements of ncAA incorporation efficiency and fidelity in BY4741 and six single-gene knockouts of BY4741 using the Alt-TAG BFP-GFP reporter in a pRS416 (URA3 marker) plasmid backbone. (B) Selected measurements of ncAA incorporation efficiency and fidelity using the Alt-TAG BFP-GFP reporter, BXG BFP-GFP reporter, and drop-in yeast display reporter in pRS416 (URA marker) plasmid backbones. The complete set of measurements using all three reporters for each of the six knockout strains strains is available in Fig. S13. (C) Architecture of drop-in yeast display system. A quantitative comparison of drop-in and conventional yeast display is available in Fig. S12. All conditions reported here were evaluated using end point measurements in biological triplicate.

Keeping these shifts in mind, we evaluated ncAA incorporation efficiency and fidelity in the parent strain and each of the knockout strains using the BXG, Alt-TAG, and drop-in display reporters. Fig. 5A depicts the results of these experiments using the Alt-TAG reporter, which exhibits the least efficient ncAA incorporation of the three reporters used here. Trends in efficiency, misincorporation, and precision observed using the Alt-TAG reporter were highly similar to the trends observed using the other two reporters (see Fig. 5B for selected data Fig. S13 depict the complete RRE and MMF data for all reporters and all strains tested here). The largest increase in ncAA incorporation efficiency we observed here was in a strain lacking PPQ1; this was consistent among all three reporters (Fig. 5B, Fig. S13). PQ1 is a protein phosphatase that has been anecdotally noted to impact stop codon readthrough, but to the best of our knowledge, its molecular role(s) in stop codon readthrough remains unknown.^49^ We note that we observed more than a two-fold gain in efficiency with the Alt-TAG reporter when PPQ1 was deleted, in contrast to the more modest gains (roughly 50 – 60%) determined with the other two reporters. This confirms the importance of controlling reporter “permissiveness” when evaluating ncAA incorporation events. Interestingly, in strains in which incorporation efficiency appeared to increase, the gains in efficiency did not appear to come at the expense of fidelity. Maximum misincorporation frequency measurements indicated similar or reduced values in comparison to MMF values in the parent strain.

To understand why we observed these increases we conducted the same analyses as described in Section 3.2 for the data depicted in Fig. 5 (Fig. S14 – S34; see Materials and Methods for full details of analysis). These analyses indicate the critical role that normalization to reporter expression level plays in determining ncAA incorporation efficiency in different strains. Increases in both N-terminal and C-terminal signal levels of wild-type and TAG-containing reporters are evident in several knockout strains. Interestingly, this includes cases in which the presence of ncAA in the induction media increases the N-terminal signal level in cells containing a reporter with an amber codon (for example, UPF2 and PQ1 knockouts with the Alt-TAG reporter). Trends in the alteration of reporter levels are different between dual-fluorescent protein reporters and the drop-in yeast display reporter. One potential explanation for this change is that only the display reporter constructs traverse the secretory pathway, where strong folding quality control mechanisms are present.^28, 50^ Changes in reporter expression may help to explain why we did not observe an increase in ncAA incorporation efficiency in a UPF1 deletion strain, which at face value appears to contradict data presented in previous reports.^35, 36^ Reporter expression levels (as detected by the N-terminal signal levels) in the UPF1 knockout vary by over 2-fold, and these variations are not fully accounted for by evaluating changes in C-terminal signal levels only (see Fig. S17, S24, S31). Both controlling for changes in reporter expression levels and performing normalization were necessary to determine whether increases in fluorescent signal were the result of enhanced ncAA incorporation efficiency. The use of a dual detection reporter architecture in combination with the RRE analysis framework accounted for both of these sources of variability in ways that would have been challenging using a single-fluorescent protein reporter. Taken together, these data demonstrate the utility of both dual-fluorescent reporters and drop-in yeast display reporters in evaluating the effects of genomic modifications on ncAA incorporation events.

## 4 Conclusions

As applications of genetically encoded noncanonical amino acids continue to mature, characterization and improvement of ncAA incorporation systems is critical to ensuring their effective deployment. This is especially true when envisioning engineered translation systems (in cells or *in vitro*) that translate alternative genetic codes with the same efficiencies as wild-type translation apparatuses and the standard genetic code. However, even in applications where efficiency requirements for ncAA incorporation are more modest, quantitative characterizations can help determine critical information such as how ncAA incorporation with a given OTS alters expression levels of a protein of interest. The work presented here provides quantitative demonstrations of how the selection of reporter type, detection methodology, and data analysis all affect the determination of ncAA incorporation efficiency and fidelity in yeast. These findings also suggest the potentially high value of conducting similar investigations in other cells and organisms. Understanding how reporters influence measurements of ncAA insertion events is an important step in facilitating cross-study comparisons of quantitative ncAA incorporation efficiency and fidelity. Single-fluorescent protein reporters, dual-fluorescent protein reporters, and yeast display reporters all exhibit similar levels of performance and precision when measured via flow cytometry. However, we identified cases when the use of single-fluorescent protein reporters could not identify changes in reporter expression levels, rather than perceived changes in ncAA incorporation effiency, suggesting that these reporters should be used with caution.

Excitingly, in this work we validated the use of dual-fluorescent reporters and a drop-in version of a yeast display reporter for evaluating the effects of gene knockouts on ncAA incorporation and found some knockouts that appear to enhance incorporation efficiency. We also found that even subtle details of reporter design and implementation alter the performance of ncAA incorporation systems. Until these changes become predictable a priori, cross-study comparisons of ncAA incorporation system performance may prove to be challenging. One straightforward way to begin to facilitate such comparisons would be to perform reference characterizations with known ncAA incorporation systems so that quantitative properties of new ncAA incorporation systems relative to a known system can be determined. This also applies to existing systems in new cell strains, existing systems used under the control of new promoter systems or expression conditions, or additional modifications. “Benchmarking” in this way would eliminate the need to immediately standardize reporter systems, although such standardization could be valuable in the longer term. Future validation of the reporter systems used in this study with highly sensitive mass spectrometry-based methods^9, 13^ could provide additional insights into developing reliable metrics for quantifying noncanonical amino acid incorporation. In any case, fully illuminating the current landscape of ncAA incorporation systems available in different organisms would enable identification of effective combinations of engineering strategies for enhancing alternative translations of the genetic code. As we enter an era in which genomes can be rapidly constructed,^27, 51–54^ evolved,^55–57^ or expanded to include additional nucleic acid bases,^19^ quantitative reporter systems with known properties will be indispensable for pushing the generation of polypeptides with chemically diverse building blocks to new heights.

## Acknowledgements

Research reported in this publication was supported by the National Institute Of General Medical Sciences of the National Institutes of Health under Award Number R35GM133471, the Army Research Office under Award Number W911NF-16-1-0175, and Tufts startup funds (to J.A.V.). J.T.S. was supported in part by an NSF Graduate Research Fellowship (ID: 2016231237). Ming Lei was supported by funds from a 2018 Tufts Collaborates grant, PI Yoon H. Kang. The content is solely the responsibility of the authors and does not necessarily represent the official views of the National Institutes of Health, the Army Research Office, or Tufts University.

## Supporting Information

**Table S1:** Flow cytometry labeling conditions.

| Detection | Primary label (dilution) | Secondary label (dilution) |
| --- | --- | --- |
| HA tag (displaying) | Mouse anti-HA (1:500) | Goat anti-mouse Alexa Fluor 488 (1:500) |
| c-Myc tag (full length) | Chicken anti-CMYC (1:500) | Goat anti-chicken Alexa Fluor 647 (1:500) |

**Table S2:** Relative readthrough efficiency error as a percentage of the RRE magnitude for three reporter systems (yeast display, RFP-GFP dual-fluorescent protein reporter, sfGFP single-fluorescent protein reporter).

|  | Detection Method | LeuOmeRS + OmeY | LeuOmeRS + AzF | TyrOmeRS + OmeY | TyrOmeRS + AzF |
| --- | --- | --- | --- | --- | --- |
| **Yeast Display** | Flow Cytometry | 13% | 15% | 17% | 9.9% |
| **RFP/GFP** | Flow Cytometry | 5.6% | 5.3% | 19% | 19% |
| **GFP** | Flow Cytometry | 10% | 2.3% | 16% | 15% |
| **RFP/GFP** | Plate Reader | 20% | 40% | 20% | 21% |
| **GFP** | Plate Reader | 19% | 140% | 22% | 19% |

**Table S3:** Relative readthrough efficiency error as a percentage of the RRE magnitude for various dual-fluorescent protein reporters.

|  | Detection Method | LeuOmeRS + OmeY | LeuOmeRS + AzF | TyrOmeRS + OmeY | TyrOmeRS + AzF |
| --- | --- | --- | --- | --- | --- |
| **RFP/GFP** | Flow Cytometry | 5.6% | 5.3% | 19% | 19% |
| **BFP/GFP** | Flow Cytometry | 15% | 13% | 7.6% | 7.5% |
| **GFP/BFP** | Flow Cytometry | 43% | 40% | 32% | 28% |
| **RFP/GFP** | Plate Reader | 20% | 40% | 20% | 21% |
| **BFP/GFP** | Plate Reader | 21% | 24% | 23% | 16% |
| **GFP/BFP** | Plate Reader | 37% | 210% | 76% | 73% |

**Fig. S1:**
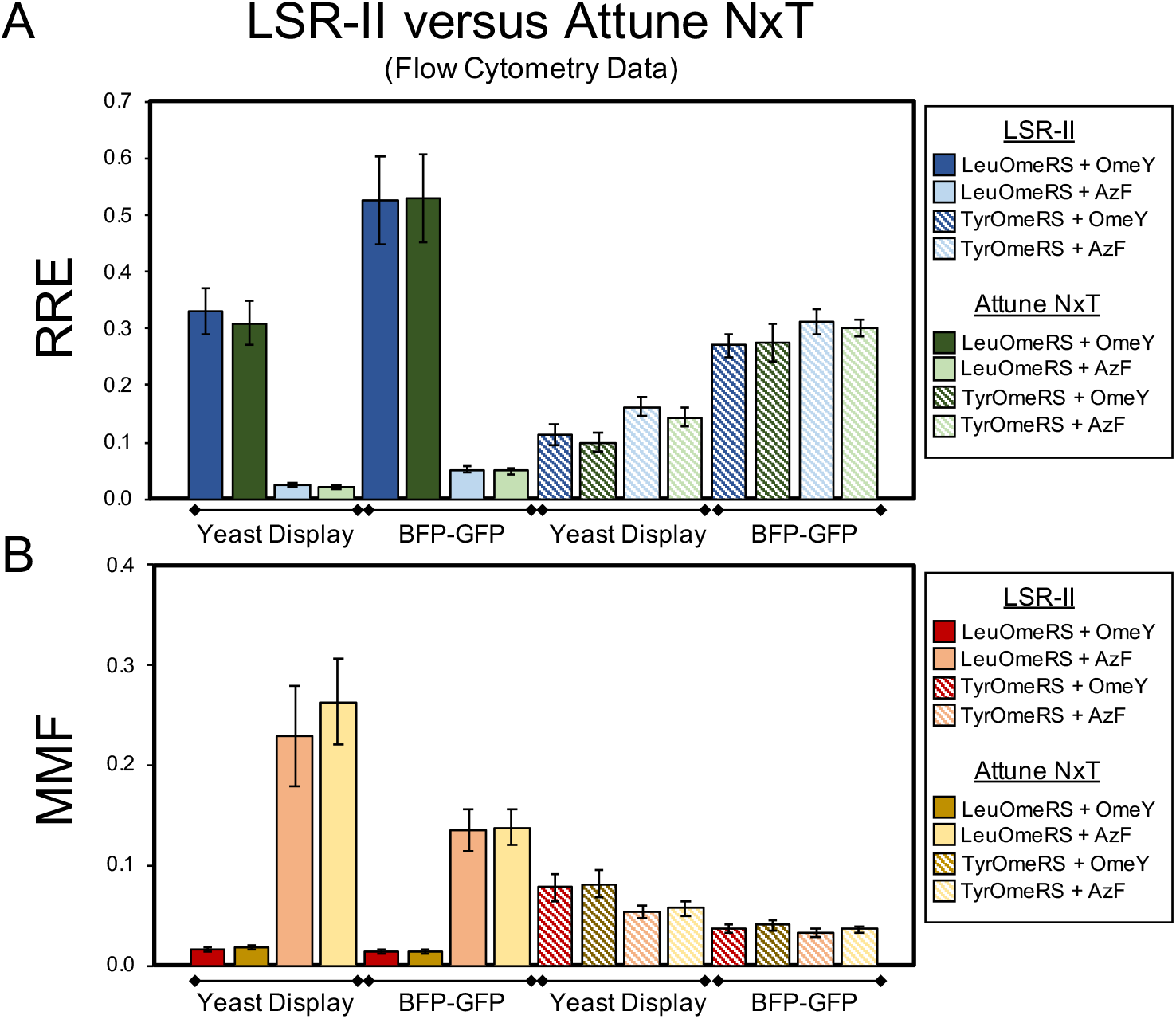
Comparison of measurements of ncAA incorporation between two different flow cytometers (LSR-II and Attune NxT). Yeast display and BFP-GFP fluorescent reporter calculations of (A) relative readthrough efficiency (RRE) and (B) maximum misincorporation frequency (MMF) with 4 orthogonal translation system-ncAA combinations run on both the LSR-II and Attune NxT. Cells from the same induced cultures were run on each instrument in these experiments.

**Fig. S2:**
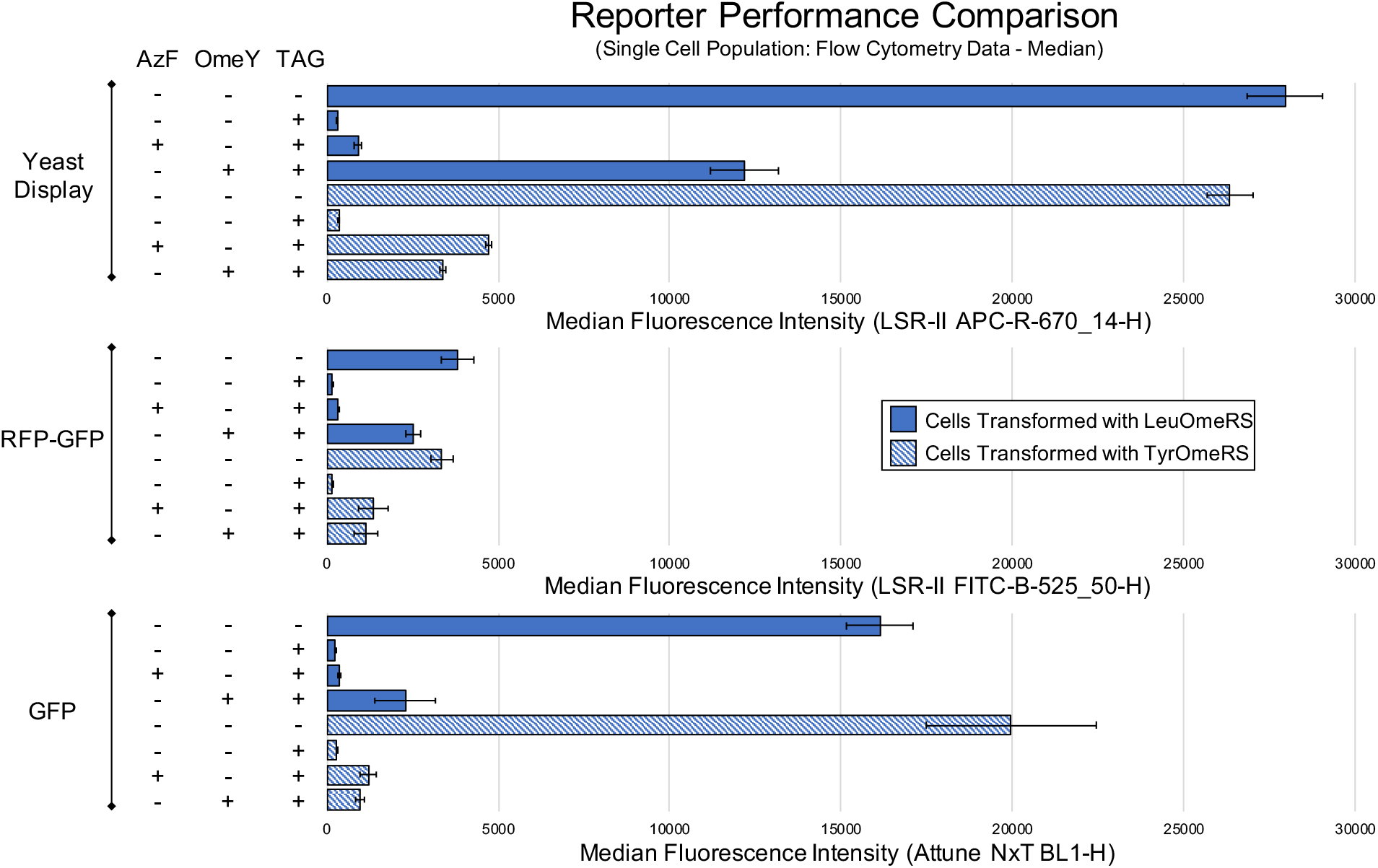
Reporter performance comparison. Quantification of ncAA incorporation efficiency for four orthogonal translation system-ncAA combinations performed with yeast display, a RFP-GFP dual-fluorescent protein reporter, and a sfGFP single-fluorescent protein reporter. Performance was evaluated with an alternative analysis method using the median fluorescence intensity of the gated single cell population (see Materials and Methods for details). The condition denoted by absence of ncAAs and a TAG codon is the wild-type reporter construct induced in the absence of ncAAs.

**Fig. S3:**
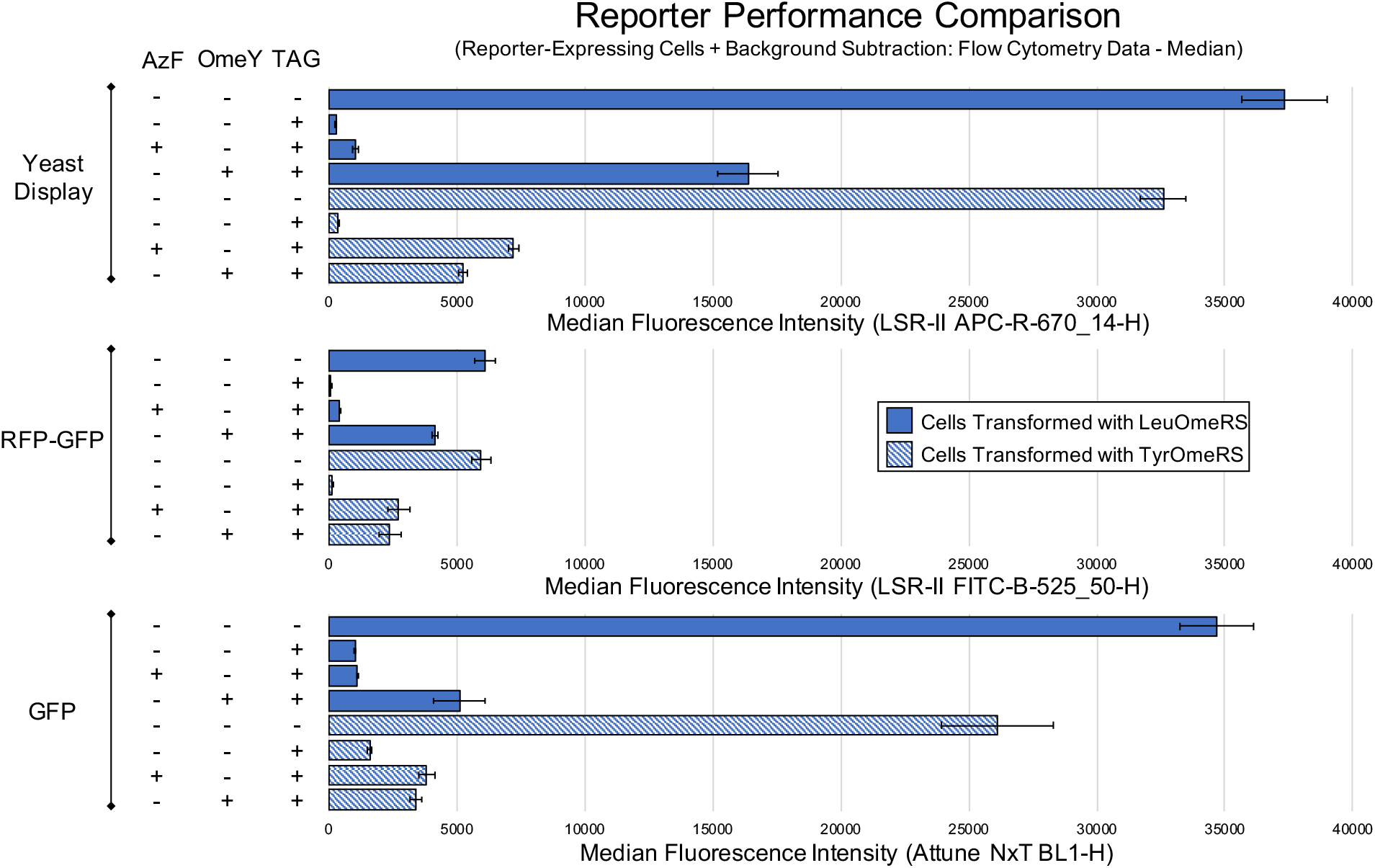
Reporter performance comparison. Quantification of ncAA incorporation efficiency for four orthogonal translation system-ncAA combinations performed with yeast display, a RFP-GFP dual-fluorescent protein reporter, and a sfGFP single-fluorescent protein reporter. Performance was evaluated with an alternative analysis method using the median fluorescence intensity of reporter-expressing cells with background subtraction. The condition denoted by absence of ncAAs and a TAG codon is the wild-type reporter construct induced in the absence of ncAAs.

**Fig. S4:**
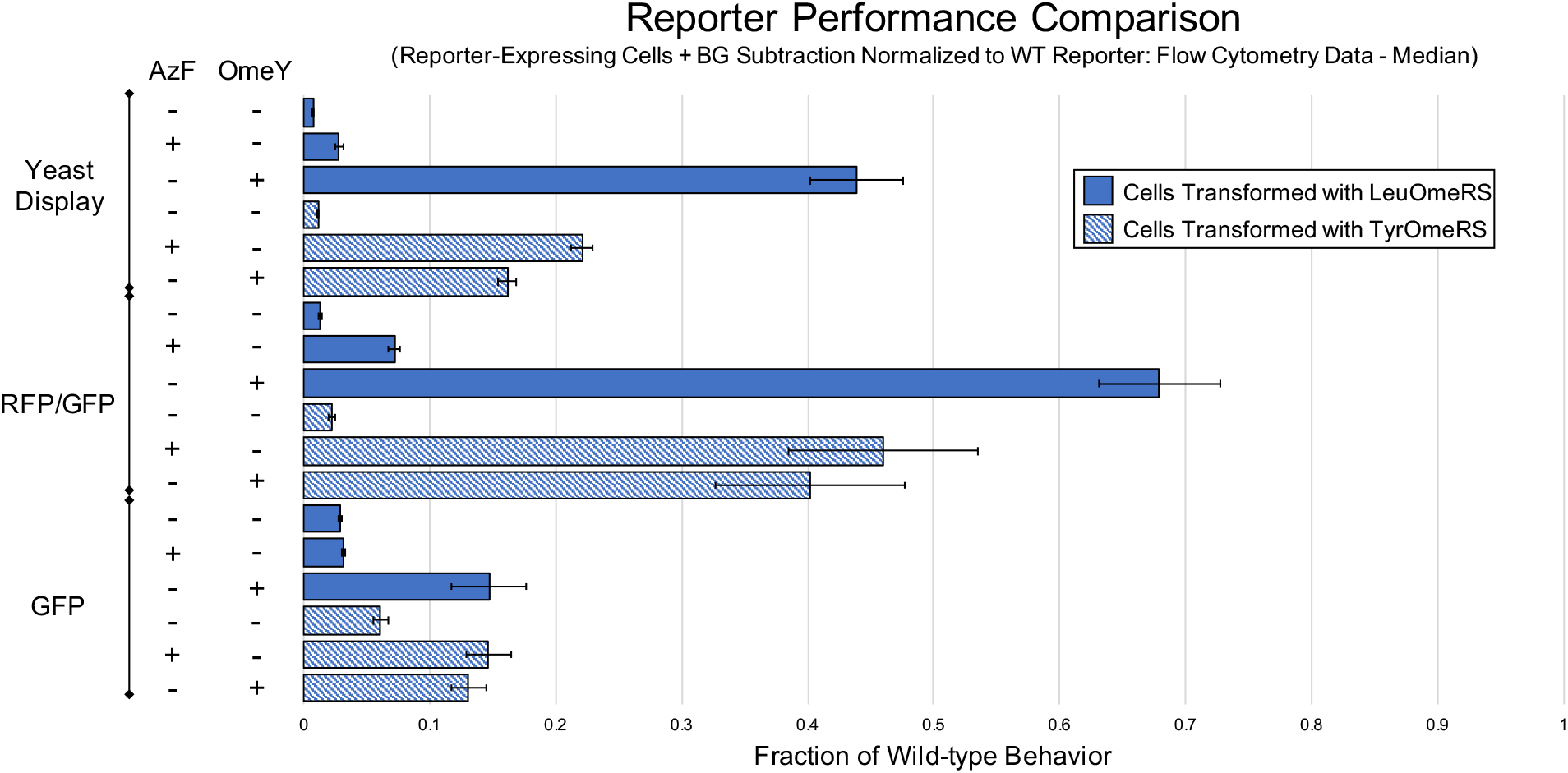
Reporter performance comparison. Quantification of ncAA incorporation efficiency for four orthogonal translation system-ncAA combinations performed with yeast display, a RFP-GFP dual-fluorescent protein reporter, and a sfGFP single-fluorescent protein reporter. Performance was evaluated with an alternative analysis method using the median fluorescence intensity of reporter-expressing cells with background subtraction and normalization to the median fluorescence intensity of the cells expressing the wild-type construct induced in the absence of ncAAs (see Materials and Methods for details).

**Fig. S5:**
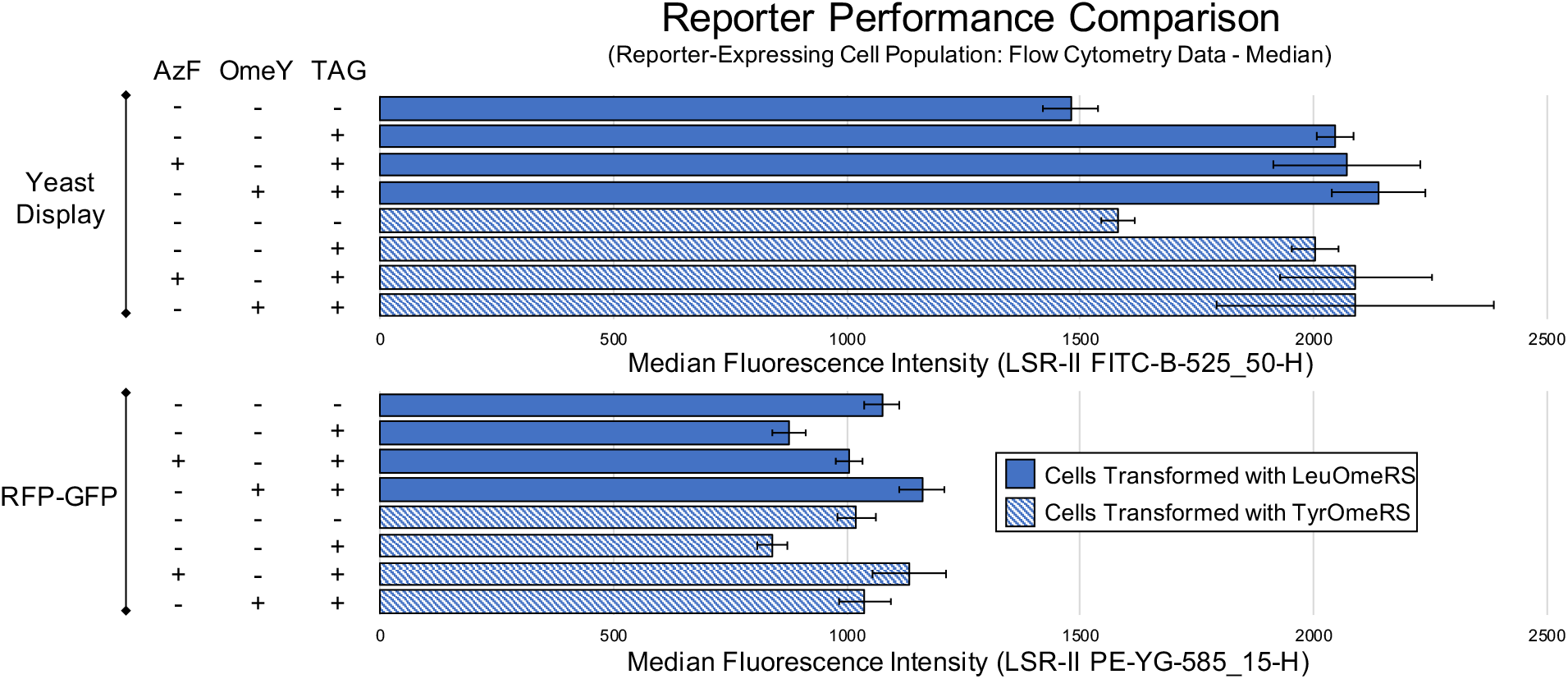
Reporter performance comparison. Measurement of reporter expression level (either yeast display or a RFP-GFP dual-fluorescent protein reporter) with an alternative analysis method using the median fluorescence intensity of the cell population exhibiting above-background reporter expression. The condition denoted by absence of ncAAs and a TAG codon is the wild-type reporter construct induced in the absence of ncAAs.

**Fig. S6:**
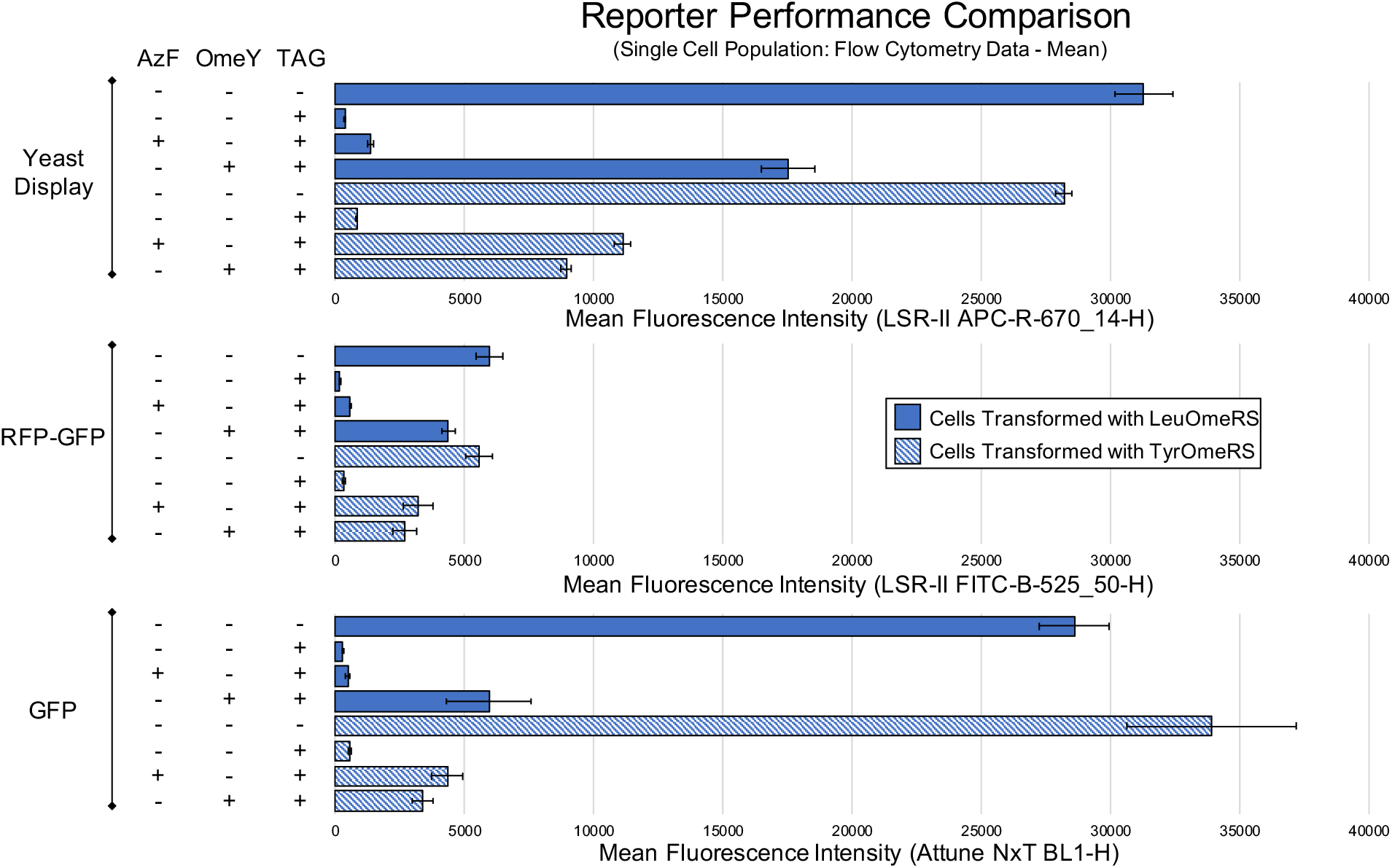
Reporter performance comparison. Quantification of ncAA incorporation efficiency for four orthogonal translation system-ncAA combinations performed with yeast display, a RFP-GFP dual-fluorescent protein reporter, and a sfGFP single-fluorescent protein reporter. Performance was evaluated with an alternative analysis method using the mean fluorescence intensity of the single cell population. The condition denoted by absence of ncAAs and a TAG codon is the wild-type reporter construct induced in the absence of ncAAs.

**Fig. S7:**
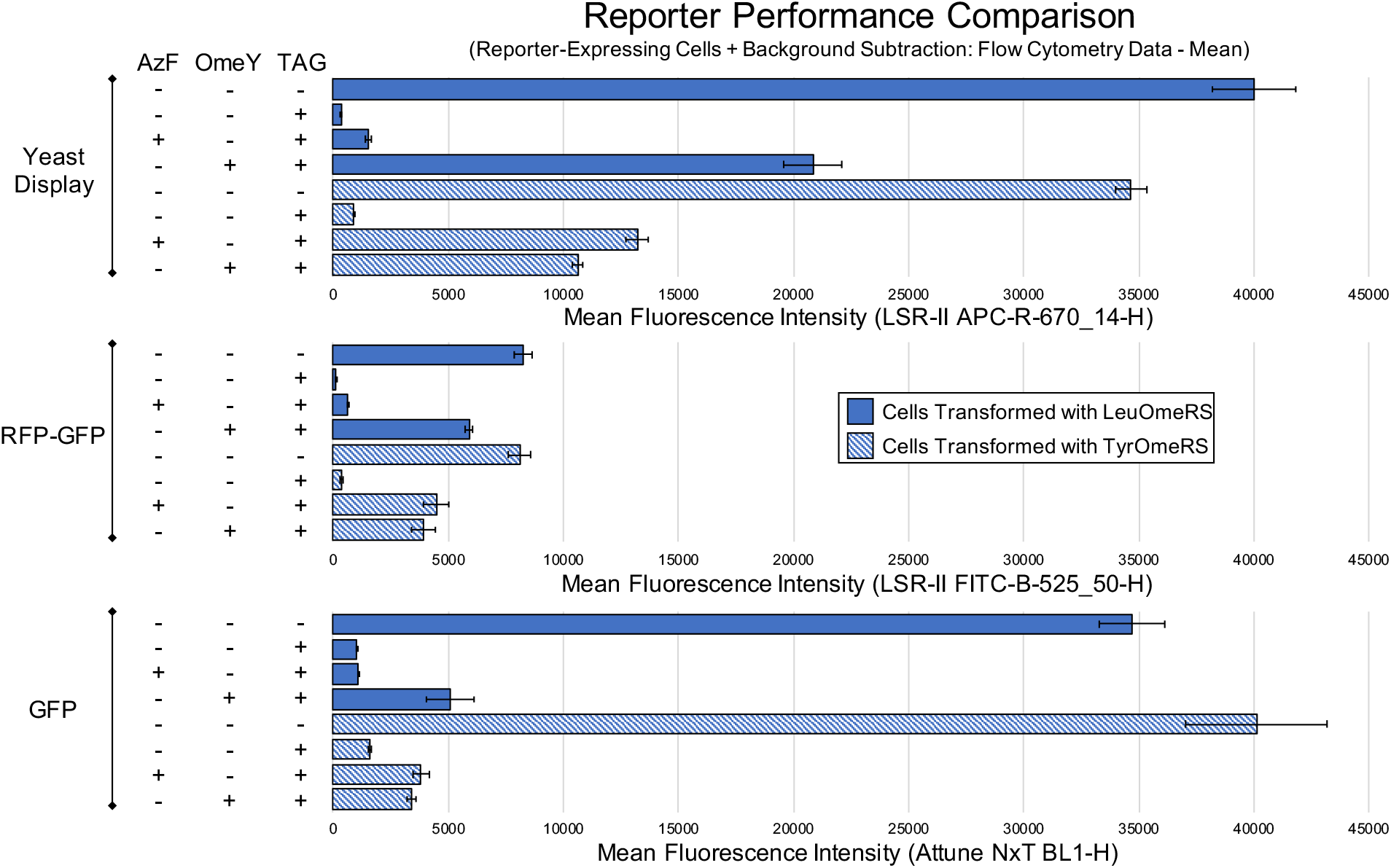
Reporter performance comparison. Quantification of ncAA incorporation efficiency for four orthogonal translation system-ncAA combinations performed with yeast display, a RFP-GFP dual-fluorescent protein reporter, and a sfGFP single-fluorescent protein reporter. Performance was evaluated with an alternative analysis method using the mean fluorescence intensity of reporter-expressing cells with background subtraction. The condition denoted by absence of ncAAs and a TAG codon is the wild-type reporter construct induced in the absence of ncAAs.

**Fig. S8:**
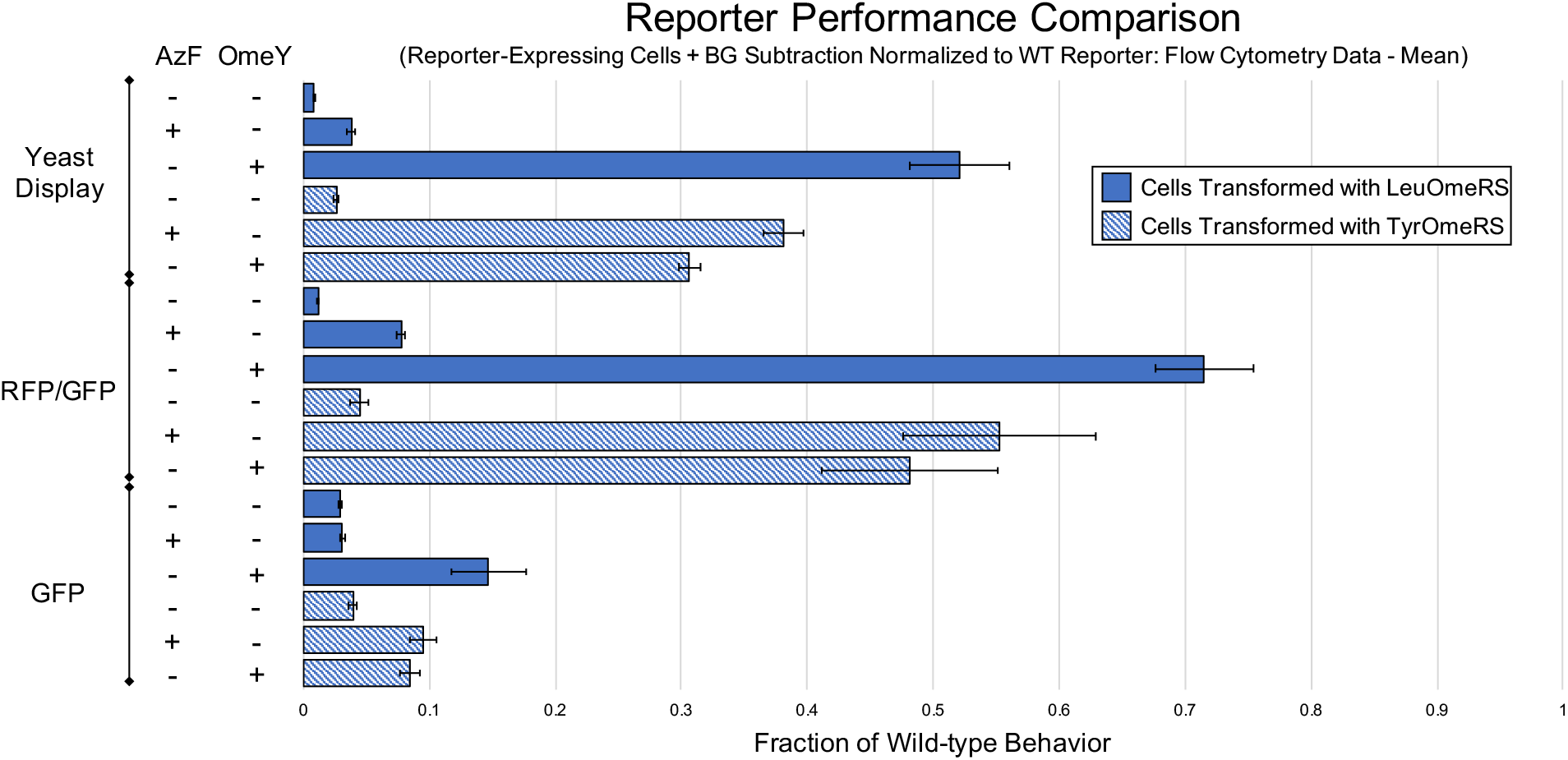
Reporter performance comparison. Quantification of ncAA incorporation efficiency for four orthogonal translation system-ncAA combinations performed with yeast display, a RFP-GFP dual-fluorescent protein reporter, and a sfGFP single-fluorescent protein reporter. Performance was evaluated with an alternative analysis method using the mean fluorescence intensity of reporter-expressing cells with background subtraction and normalized to the mean fluorescence intensity of the cells expressing the wild-type construct induced in the absence of ncAAs.

**Fig. S9:**
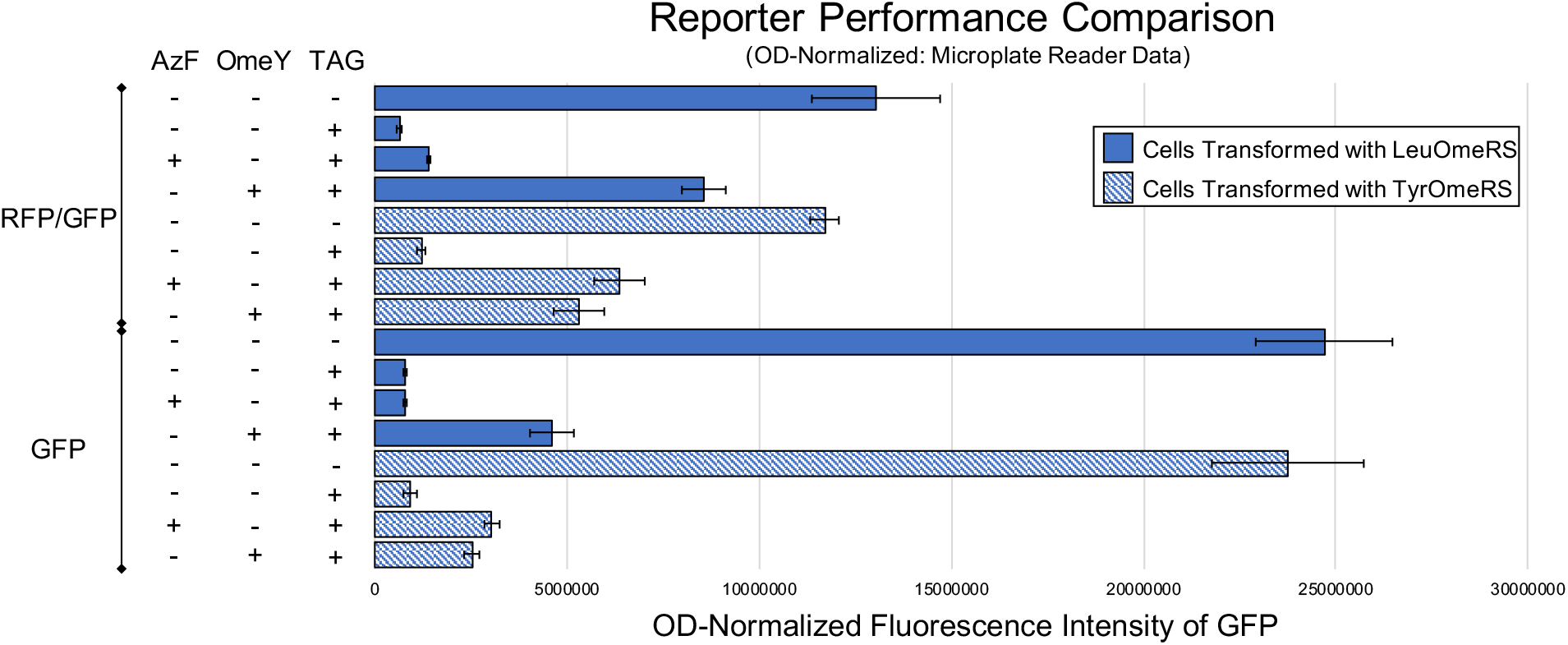
Reporter performance comparison. Quantification of ncAA incorporation efficiency for four orthogonal translation system-ncAA combinations performed with a RFP-GFP dual-fluorescent protein reporter, and a sfGFP single-fluorescent protein reporter and analyzed on a microplate reader. Performance was evaluated with an alternative analysis method using sample fluorescence normalized to sample OD600. The condition denoted by absence of ncAAs and a TAG codon is the wild-type reporter construct induced in the absence of ncAAs.

**Fig. S10:**
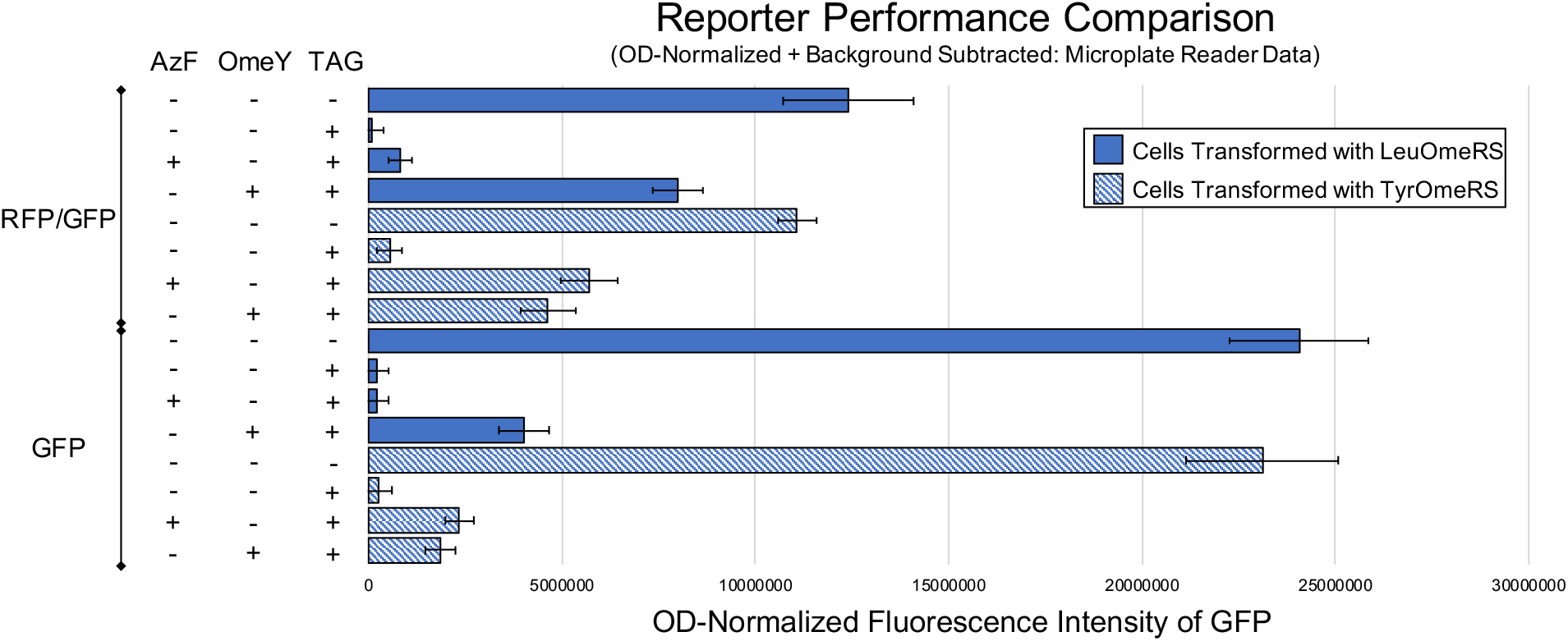
Reporter performance comparison. Quantification of ncAA incorporation efficiency for four orthogonal translation system-ncAA combinations performed with a RFP-GFP dual-fluorescent protein reporter, and a sfGFP single-fluorescent protein reporter and analyzed on a microplate reader. Performance was evaluated with an alternative analysis method using sample fluorescence normalized to sample OD600 and followed by background subtraction to account for cellular autofluorescence. The condition denoted by absence of ncAAs and a TAG codon is the wild-type reporter construct induced in the absence of ncAAs.

**Fig. S11:**
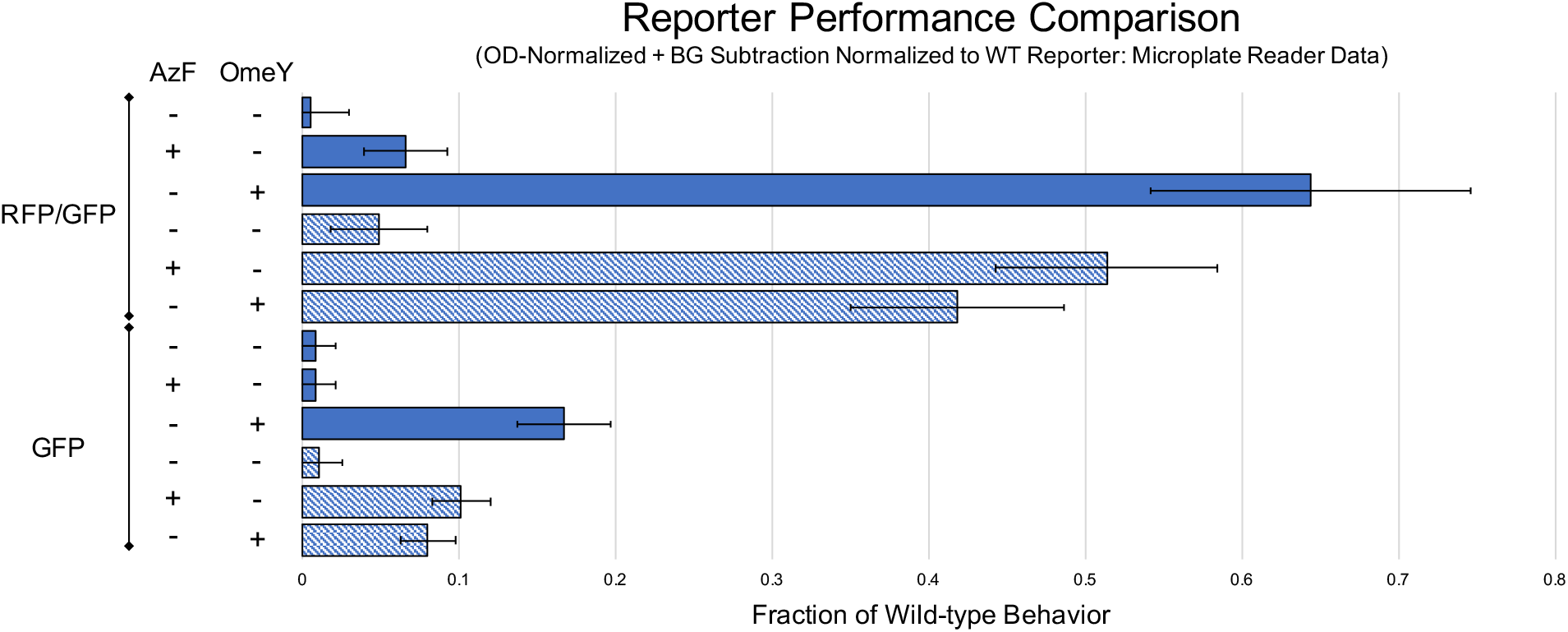
Reporter performance comparison. Quantification of ncAA incorporation efficiency for four orthogonal translation system-ncAA combinations performed with a RFP-GFP dual-fluorescent reporter, and a sfGFP single-fluorescent reporter. Performance was evaluated with an alternative analysis method using sample fluorescence normalized to sample OD600 with background subtraction and normalization to the cells expressing the wild-type reporter (also OD600 normalized) induced in the absence of ncAAs.

**Figure S12:**
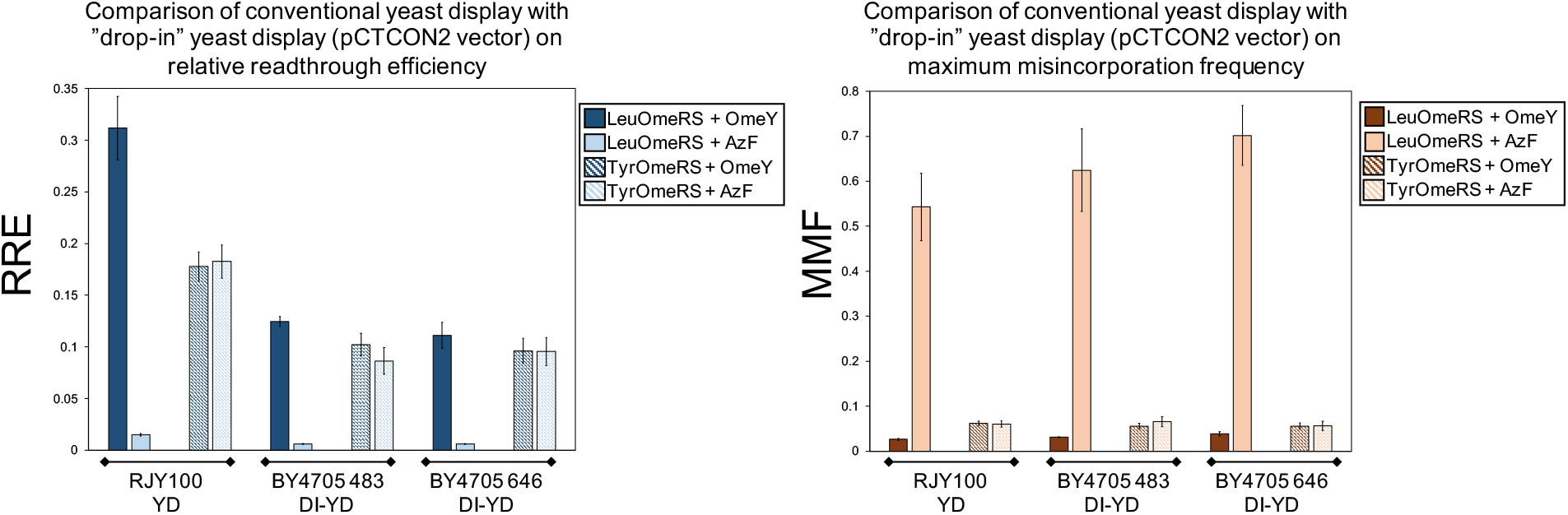
Conventional yeast display (YD) versus drop-in yeast display (DI-YD). Quantification of ncAA incorporation efficiency and fidelity for conventional display (in RJY100) and drop-in display (in multiple ΔTRP1 strains) in a pCTCON2 vector backbone.

**Figure S13:**
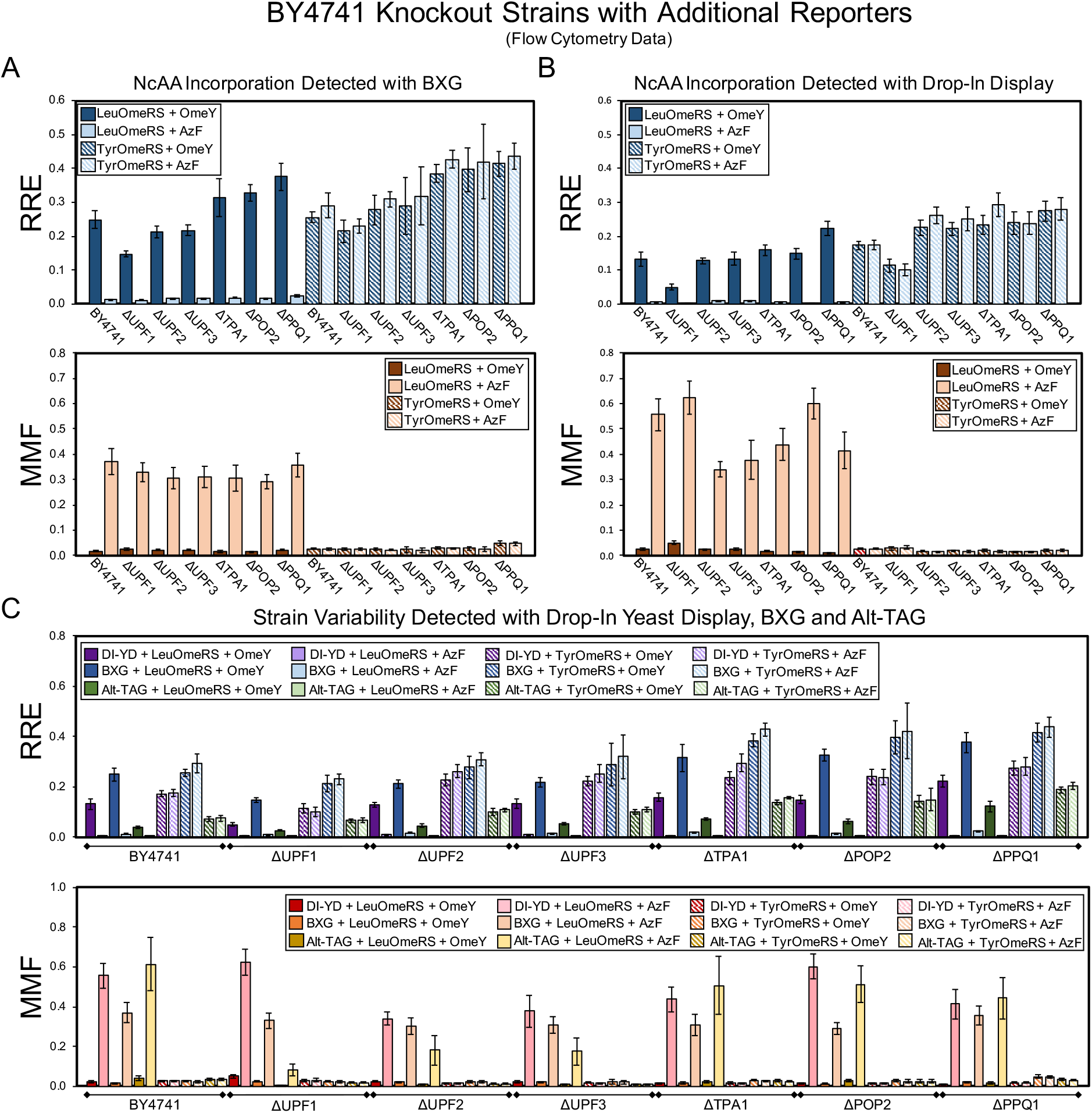
Evaluation of ncAA incorporation events in a series of single-gene knockout strains using two dual-fluorescent protein reporters and a drop-in yeast display (DI-YD) reporter. (A) Measurements of ncAA incorporation efficiency and fidelity in BY4741 and six single-gene knockouts of BY4741 using the BXG BFP-GFP reporter in a pRS416 (URA3 marker) plasmid backbone. (B) Measurements of ncAA incorporation efficiency and fidelity in BY4741 and six single-gene knockouts of BY4741 using the drop-in yeast display reporter in a pRS416 (URA3 marker) plasmid backbone. (C) Complete set of ncAA incorporation measurements for efficiency and fidelity using the Alt-TAG BFP-GFP reporter, BXG BFP-GFP reporter, and drop-in yeast display reporter in pRS416 (URA marker) plasmid backbones. All conditions reported here were evaluated with end point measurements in biological triplicate.

**Fig. S14:**
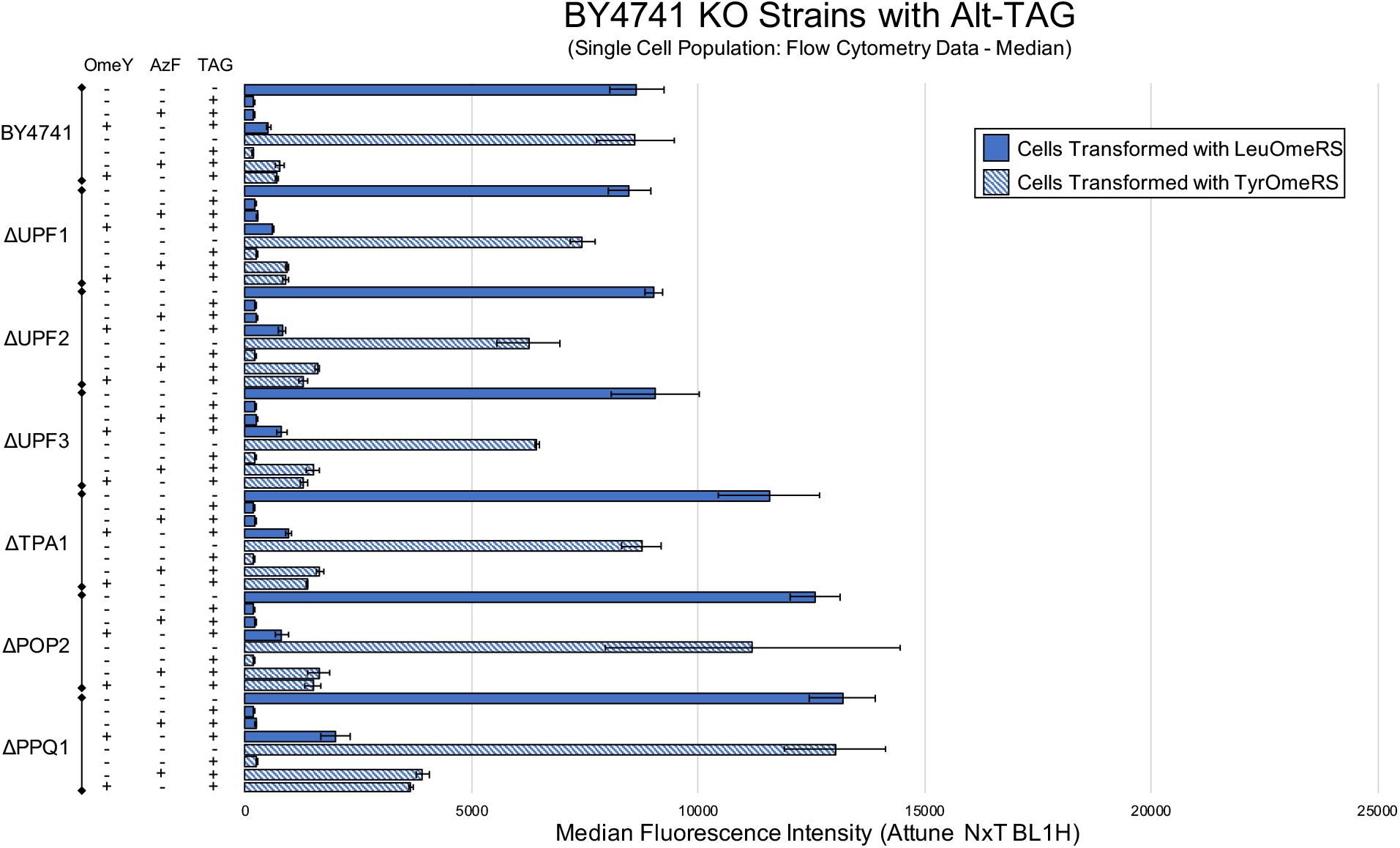
Alternate analysis of BY4741 and six single-gene knockout strains using the Alt-TAG BFP-GFP dual-fluorescent protein reporter. Measurements for the efficiency of ncAA incorporation calculated with median fluorescence intensity using single cell population analysis as described in Materials and Methods Section 2.11. The condition denoted by absence of ncAAs and a TAG codon is the wild-type reporter construct induced in the absence of ncAAs.

**Fig. S15:**
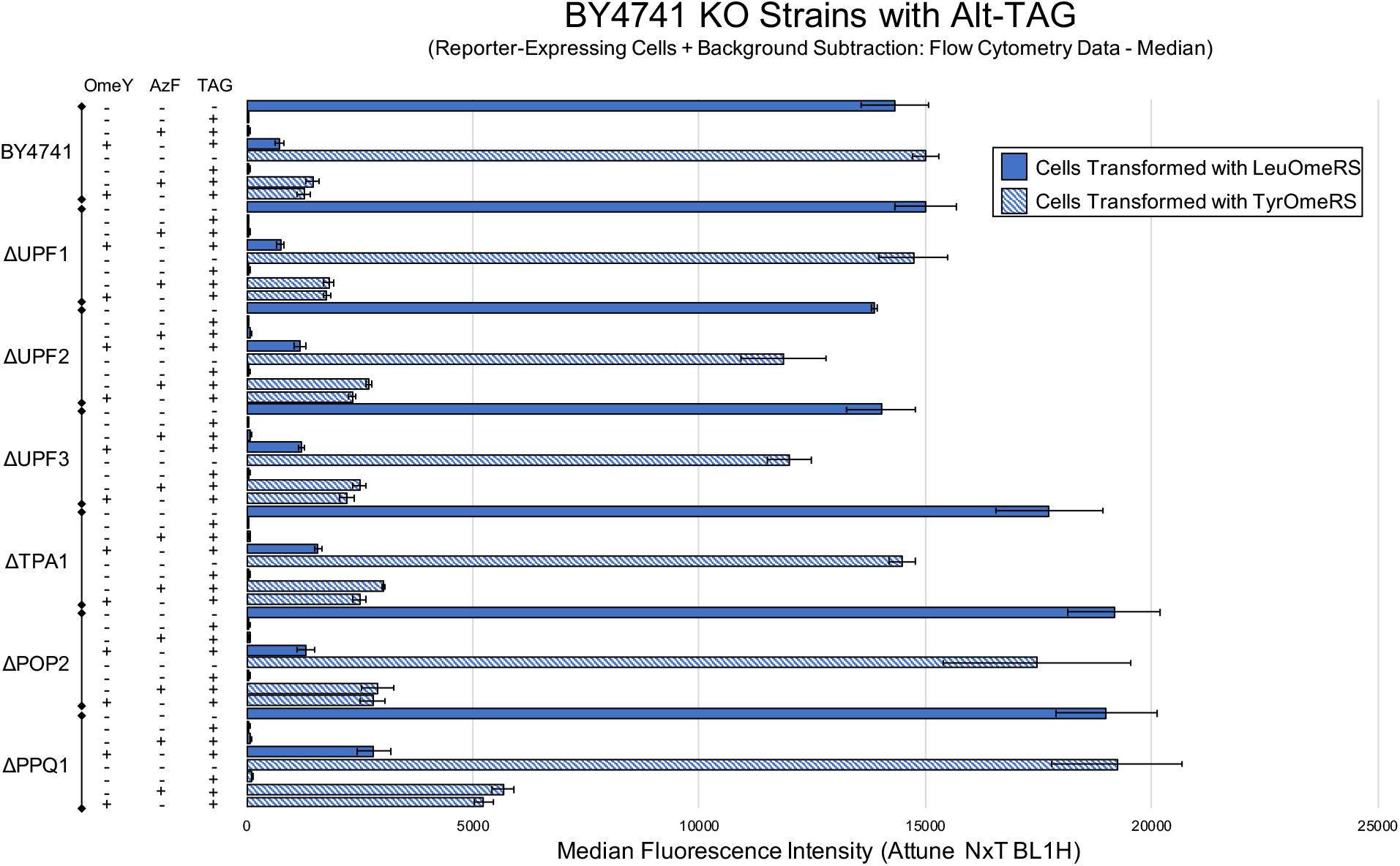
Alternate analysis of BY4741 and six single-gene knockout strains using the Alt-TAG BFP-GFP dual-fluorescent protein reporter. Measurements for the efficiency of ncAA incorporation calculated using the median fluorescence intensity of reporter-expressing cells with background subtraction as described in Materials and Methods Section 2.11. The condition denoted by absence of ncAA and TAG codon is the wild-type reporter construct induced in the absence of ncAAs.

**Fig. S16:**
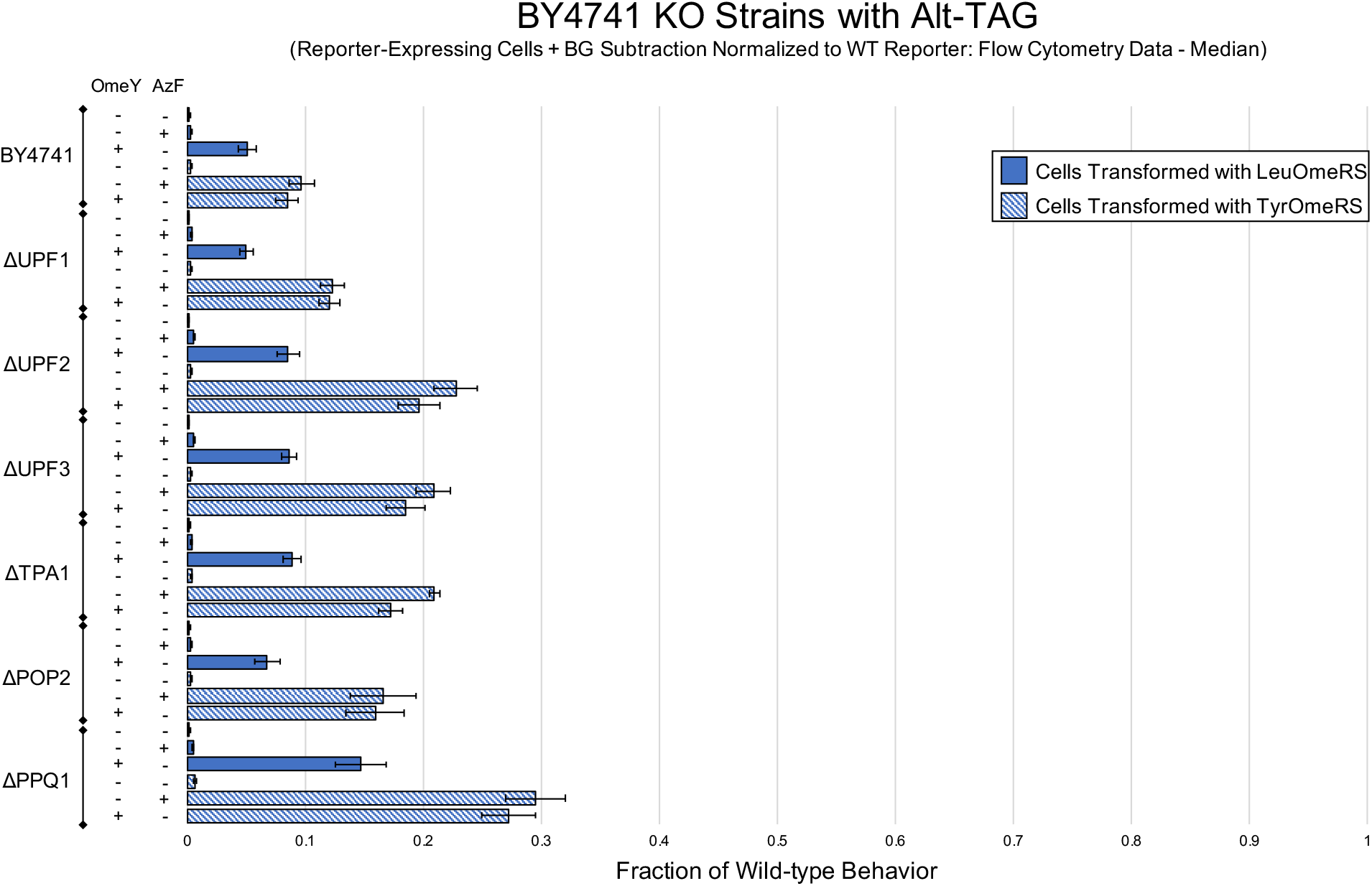
Alternate analysis of BY4741 and six single-gene knockout strains using the Alt-TAG BFP-GFP dual-fluorescent protein reporter. Measurements for the efficiency of ncAA incorporation calculated using the median fluorescence intensity of reporter-expressing cells with background subtraction and normalized to expression levels of cells expressing the wild-type reporter induced in the absence of ncAAs.

**Fig. S17:**
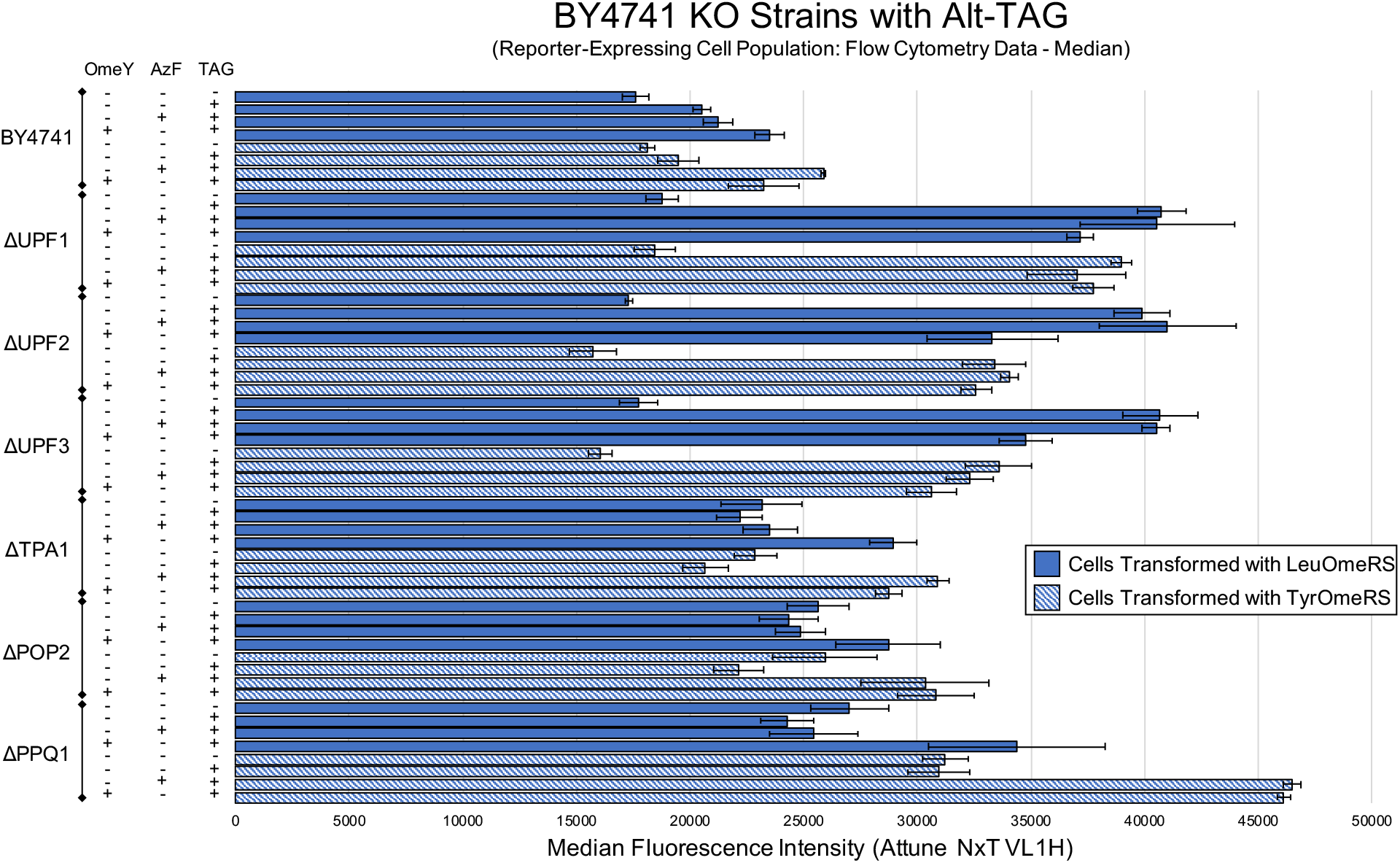
Measurement of reporter expression level in BY4741 and six single-gene knockout strains containing Alt-TAG BFP-GFP using the median fluorescence intensity of the cell population exhibiting above-background reporter expression. The condition denoted by absence of ncAAs and a TAG codon is the wild-type reporter construct induced in the absence of ncAAs.

**Fig. S18:**
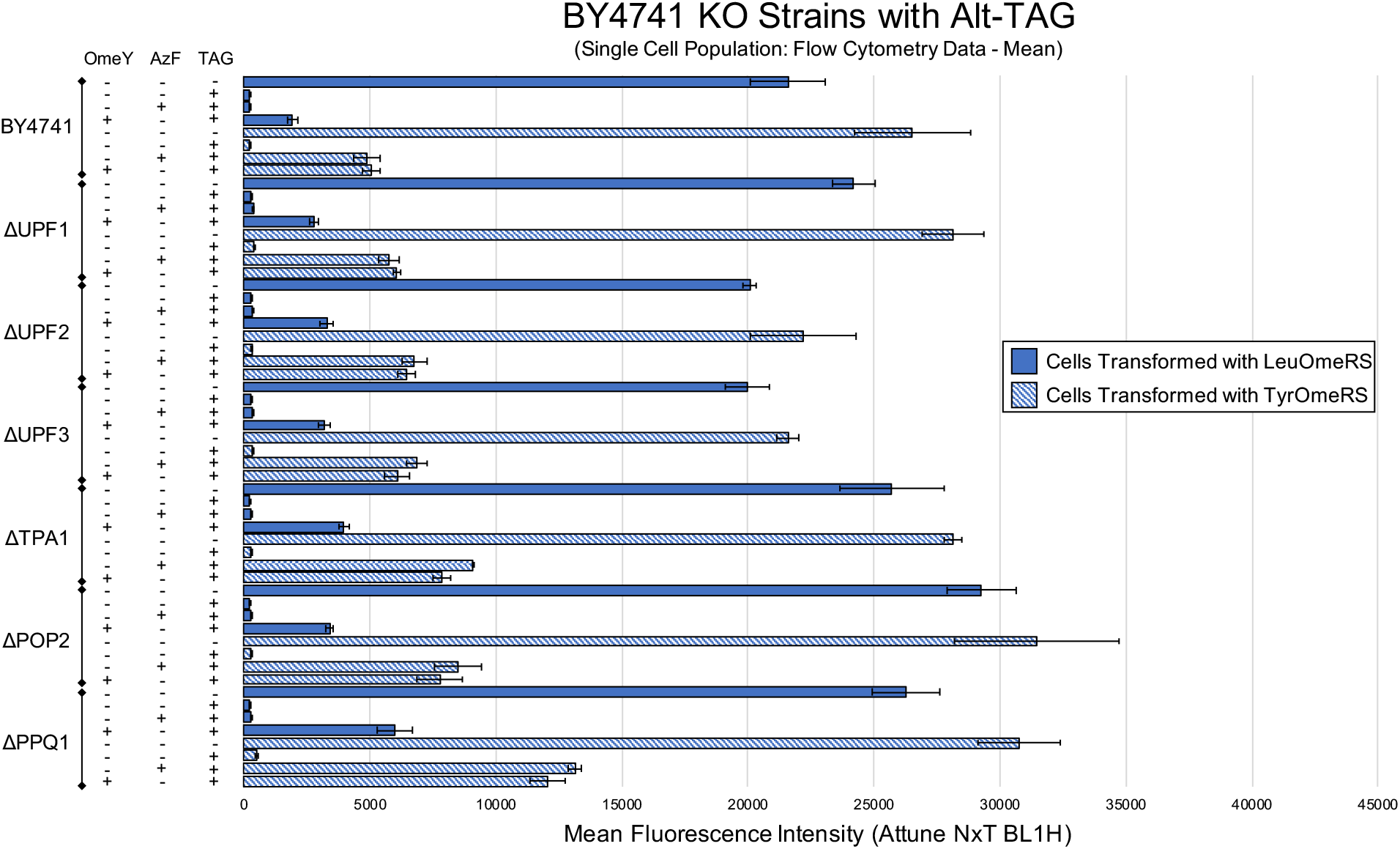
Alternate analysis of BY4741 and six single-gene knockout strains using the Alt-TAG BFP-GFP dual-fluorescent protein reporter. Performance was evaluated using the mean fluorescence intensity of the single cell population. The condition denoted by absence of ncAAs and a TAG codon is the wild-type reporter construct induced in the absence of ncAAs.

**Fig. S19:**
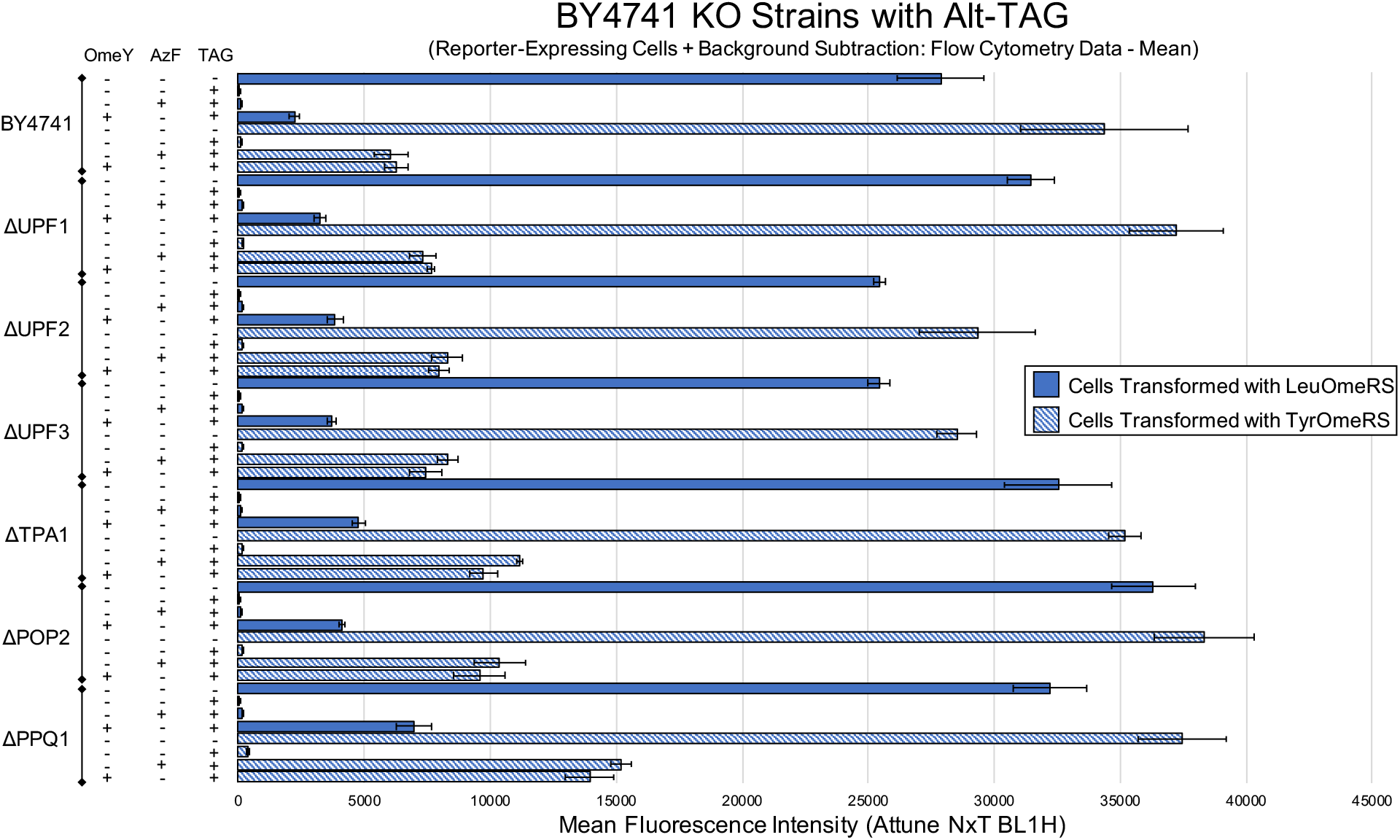
Alternate analysis of BY4741 and six single-gene knockout strains using the Alt-TAG BFP-GFP dual-fluorescent protein reporter. Performance was evaluated using the mean fluorescence intensity of reporter-expressing cells with background subtraction. The condition denoted by absence of ncAAs and a TAG codon is the wild-type reporter construct induced in the absence of ncAAs.

**Fig. S20:**
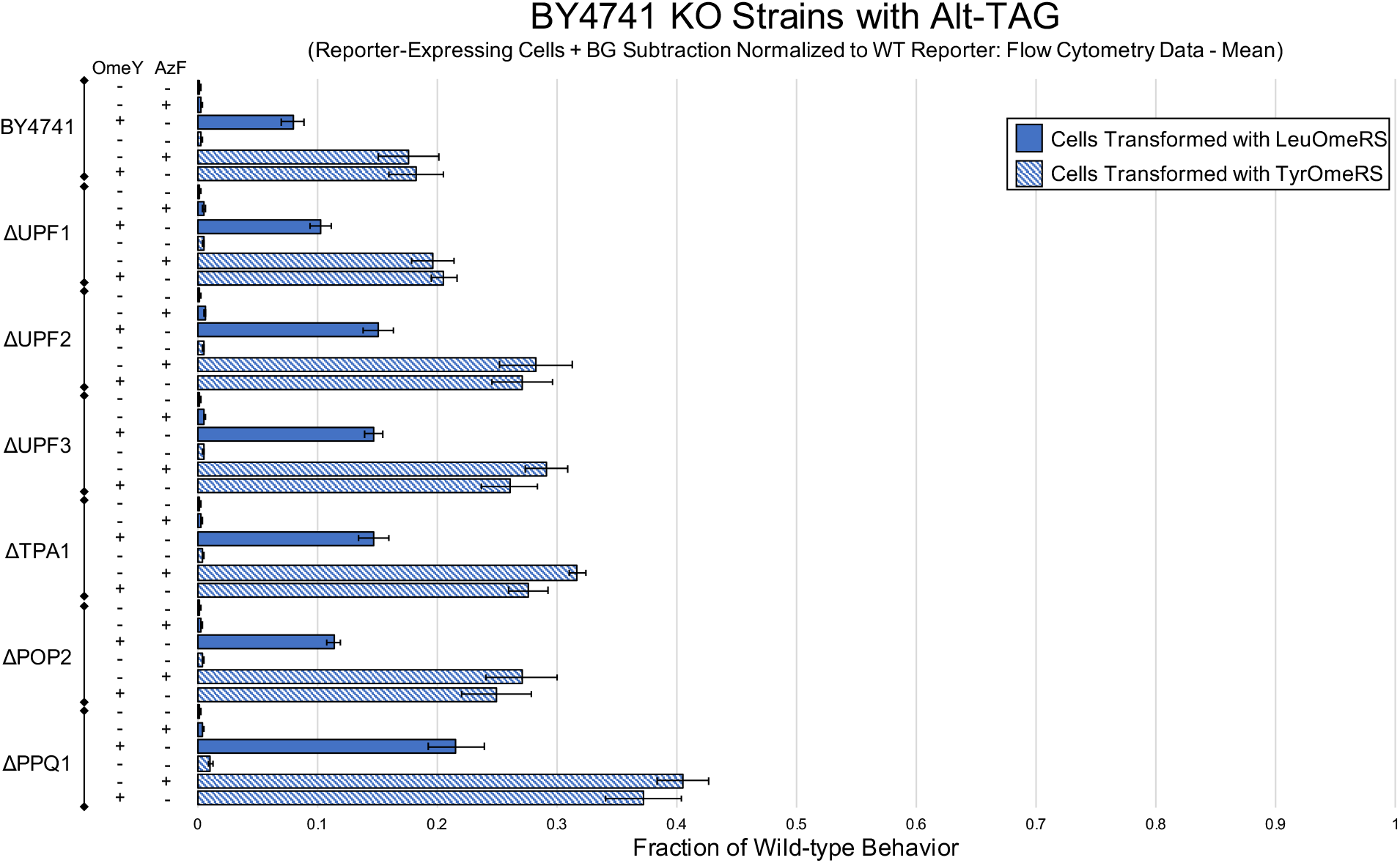
Alternate analysis of BY4741 and six single-gene knockout strains using the Alt-TAG BFP-GFP dual-fluorescent protein reporter. Measurements for the efficiency of ncAA incorporation calculated using the mean fluorescence intensity of reporter-expressing cells with background subtraction and normalized to mean expression levels of cells expressing the wild-type reporter induced in the absence of ncAAs.

**Fig. S21:**
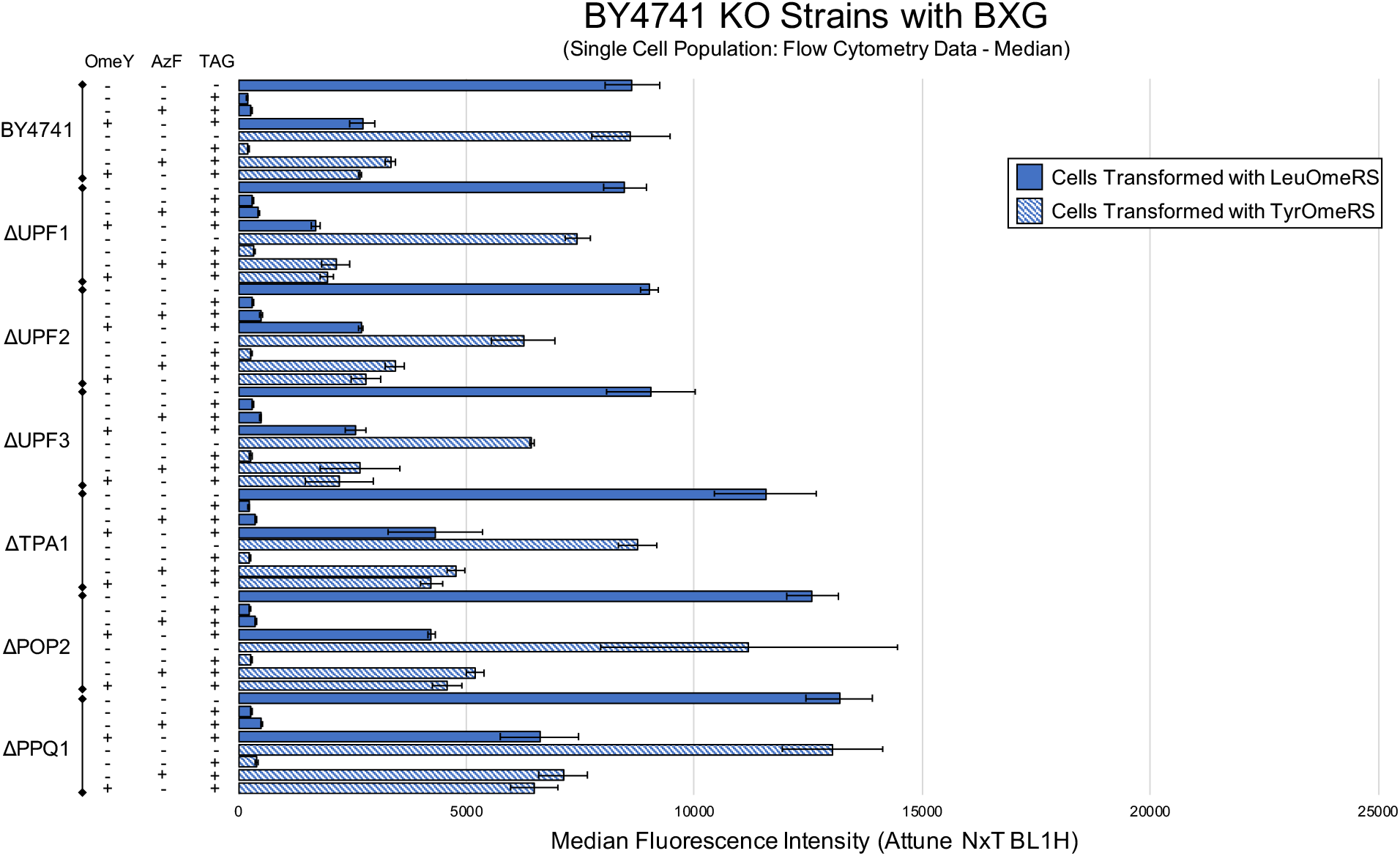
Alternate analysis of BY4741 and six single-gene knockout strains using the BXG BFP-GFP dual-fluorescent protein reporter. Measurements for the efficiency of ncAA incorporation calculated with median fluorescence intensity using single cell population analysis as described in Materials and Methods Section 2.11. The condition denoted by absence of ncAAs and a TAG codon is the wild-type reporter construct induced in the absence of ncAAs.

**Fig. S22:**
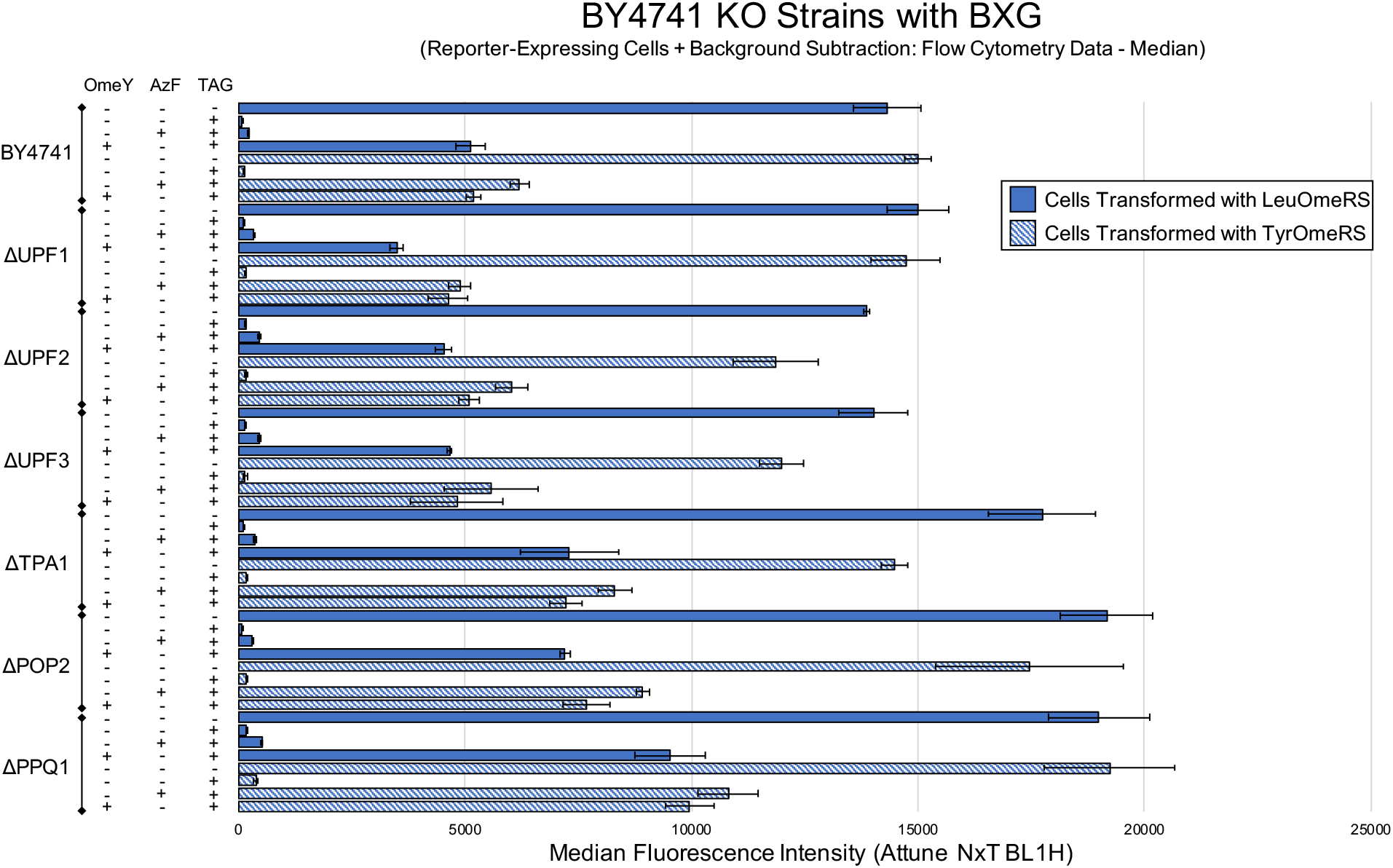
Alternate analysis of BY4741 and six single-gene knockout strains using the BXG BFP-GFP dual-fluorescent protein reporter. Measurements for the efficiency of ncAA incorporation calculated with median fluorescence intensity using reporter-expressing cells with background subtraction. The condition denoted by absence of ncAAs and a TAG codon is the wild-type reporter construct induced in the absence of ncAAs.

**Fig. S23:**
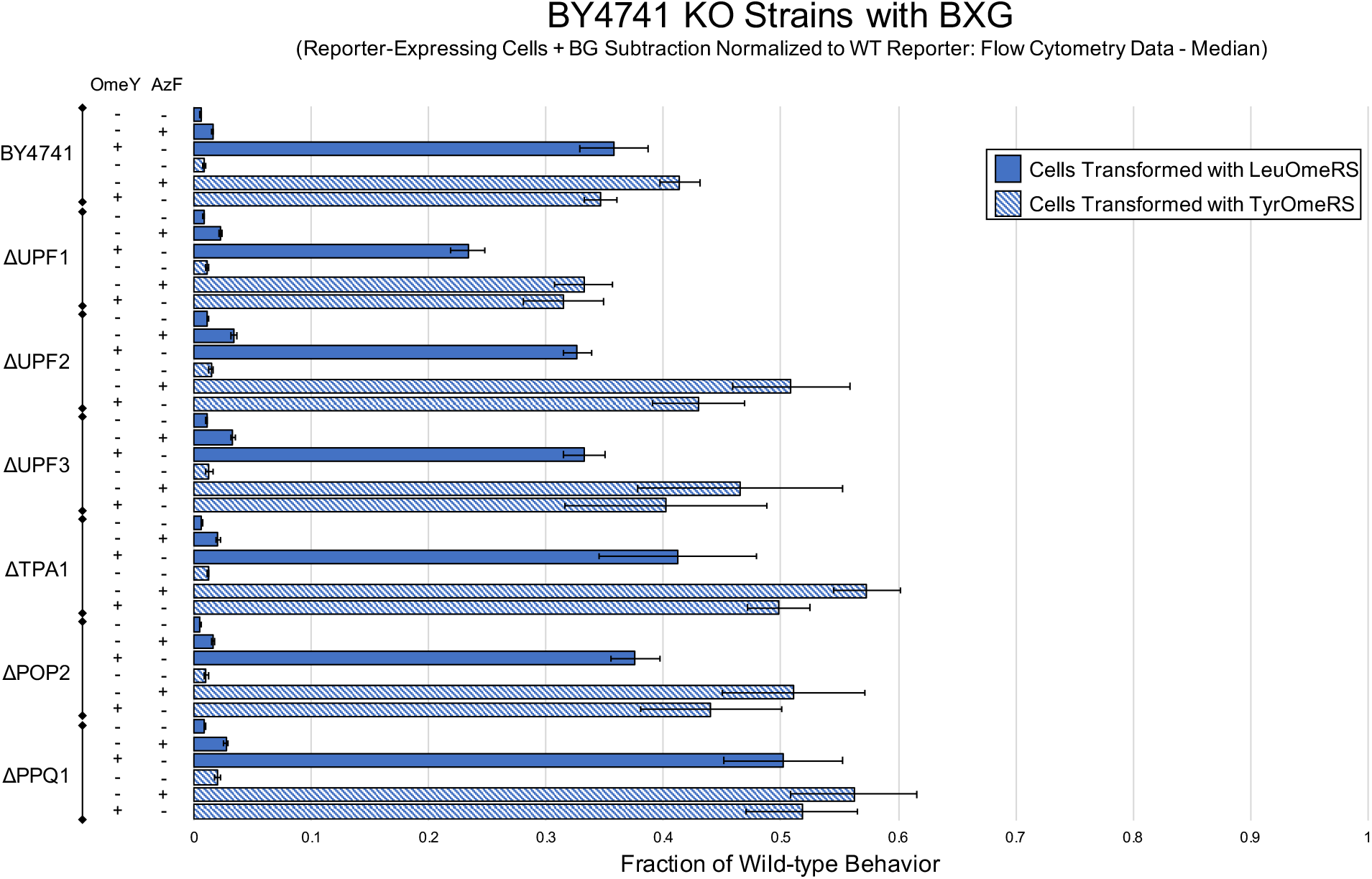
Alternate analysis of BY4741 and six single-gene knockout strains using the BXG BFP-GFP dual-fluorescent protein reporter. Measurements for the efficiency of ncAA incorporation calculated using the median fluorescence intensity of reporter-expressing cells with background subtraction and normalized to expression levels of cells expressing the wild-type reporter induced in the absence of ncAAs.

**Fig. S24:**
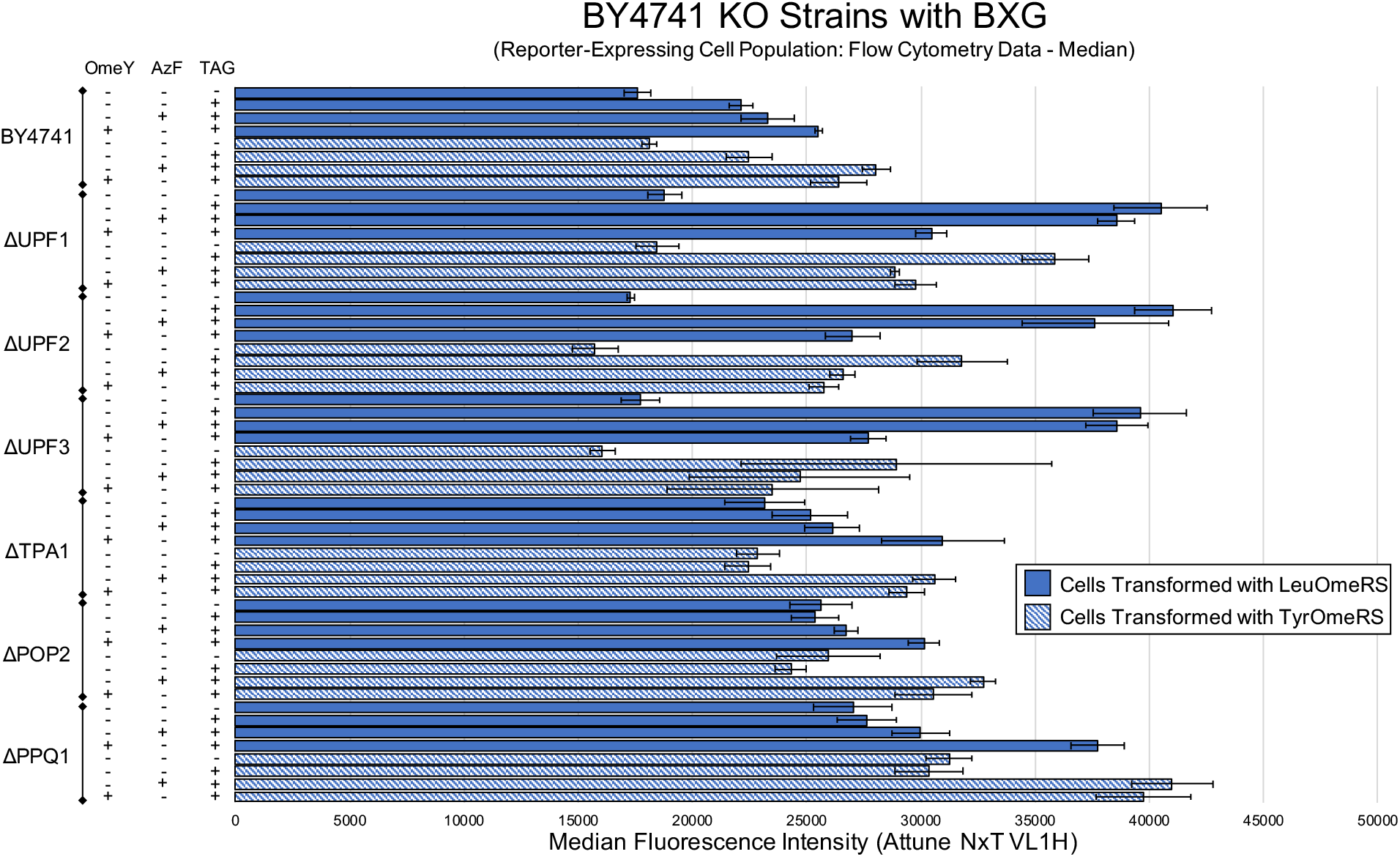
Measurement of reporter expression level in BY4741 and six single-gene knockout strains containing BXG BFP-GFP using the median fluorescence intensity of the cell population exhibiting above-background reporter expression. The condition denoted by absence of ncAAs and a TAG codon is the wild-type reporter construct induced in the absence of ncAAs.

**Fig. S25:**
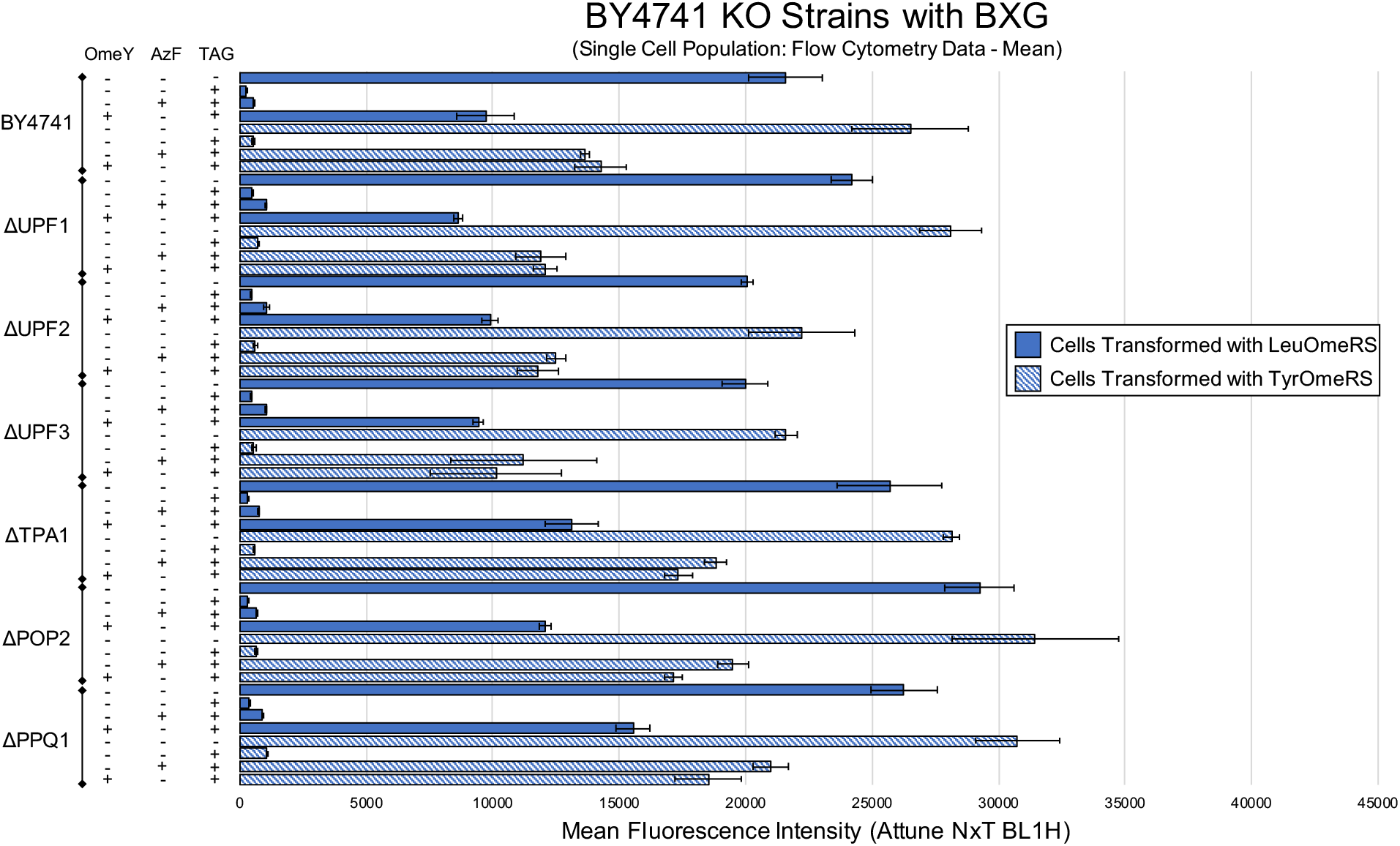
Alternate analysis of BY4741 and six single-gene knockout strains using the BXG BFP-GFP dual-fluorescent protein reporter. Measurements for the efficiency of ncAA incorporation calculated with mean fluorescence intensity using single cell population analysis as described in Materials and Methods Section 2.11. The condition denoted by absence of ncAAs and a TAG codon is the wild-type reporter construct induced in the absence of ncAAs.

**Fig. S26:**
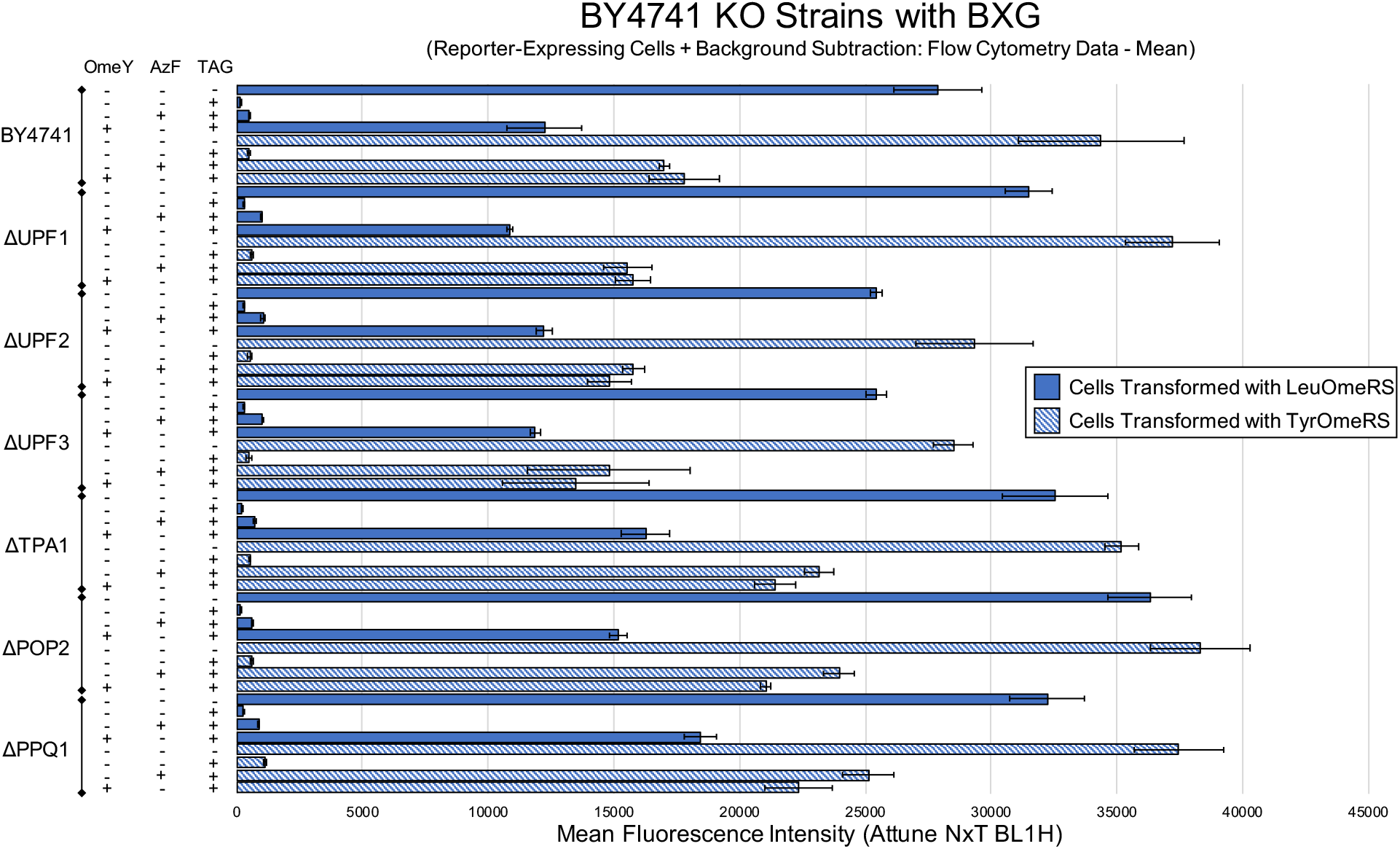
Alternate analysis of BY4741 and six single-gene knockout strains using the BXG BFP-GFP dual-fluorescent protein reporter. Measurements for the efficiency of ncAA incorporation calculated with mean fluorescence intensity using reporter-expressing cells with background subtraction. The condition denoted by absence of ncAAs and a TAG codon is the wild-type reporter construct induced in the absence of ncAAs.

**Fig. S27:**
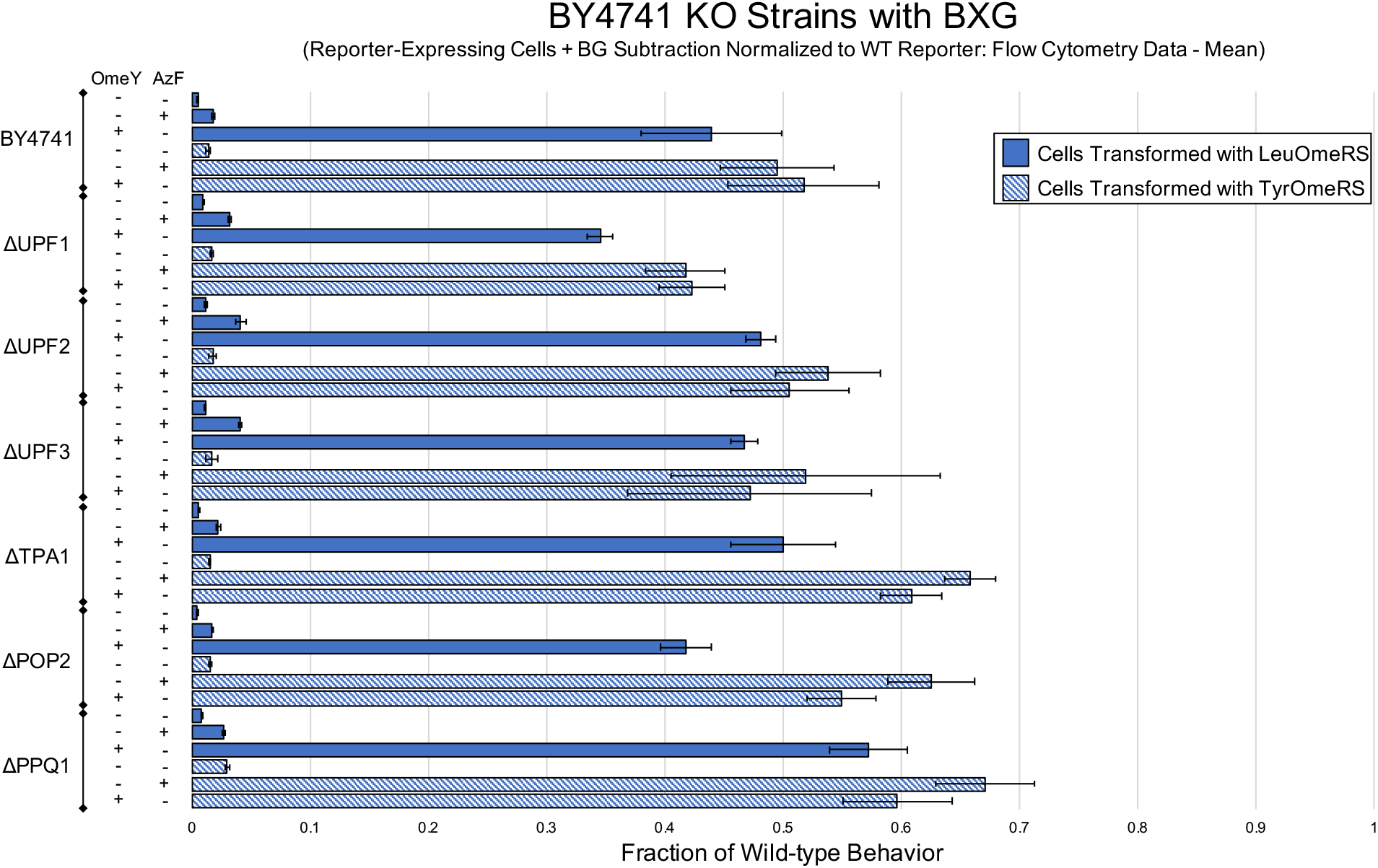
Alternate analysis of BY4741 and six single-gene knockout strains using the BXG BFP-GFP dual-fluorescent protein reporter. Measurements for the efficiency of ncAA incorporation calculated using the mean fluorescence intensity of reporter-expressing cells with background subtraction and normalized to mean expression levels of cells expressing the wild-type reporter induced in the absence of ncAAs.

**Fig. S28:**
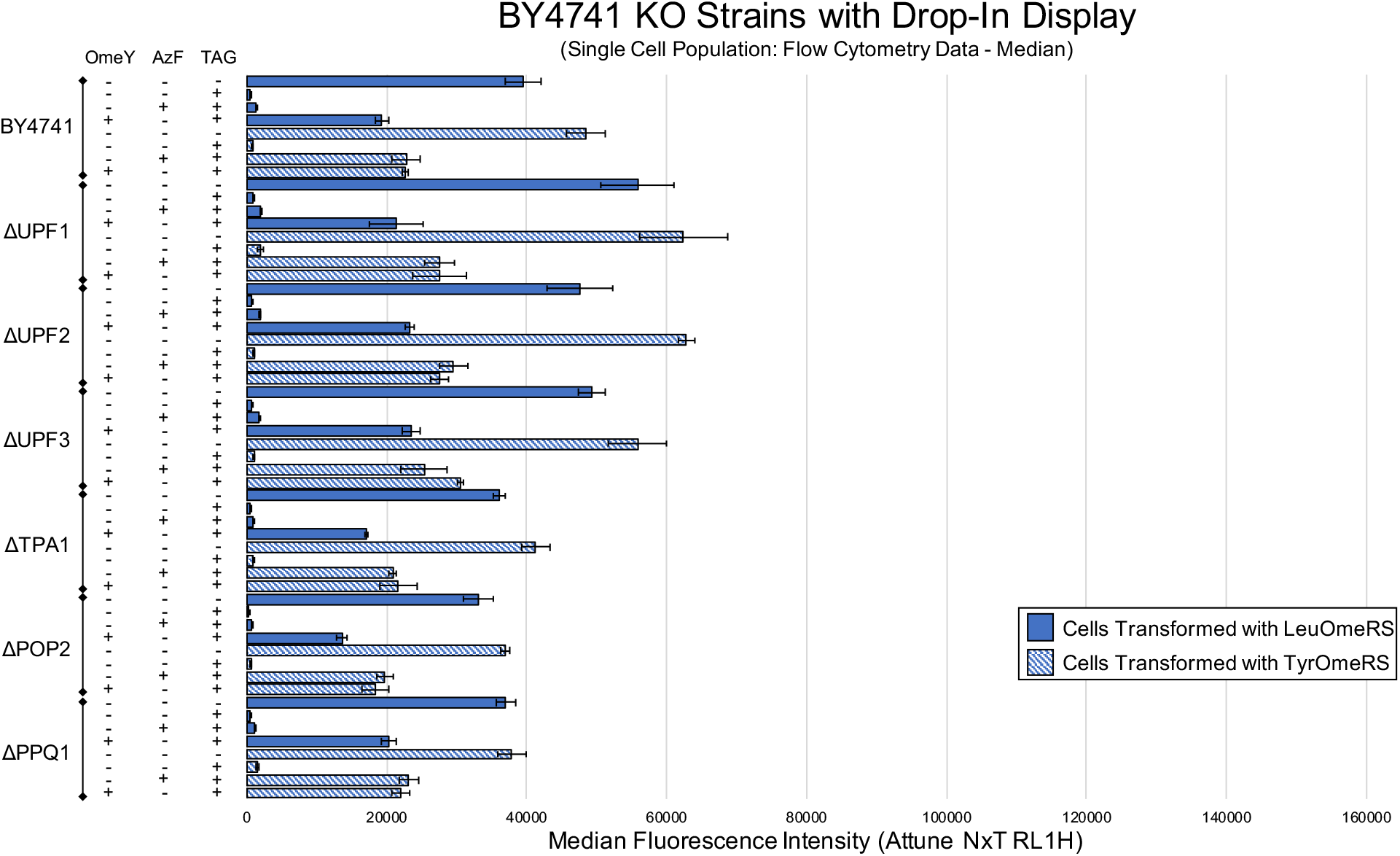
Alternate analysis of BY4741 and six single-gene knockout strains using the drop-in yeast display reporter. Measurements for the efficiency of ncAA incorporation calculated with median fluorescence intensity using single cell population analysis as described in Materials and Methods Section 2.11. The condition denoted by absence of ncAAs and a TAG codon is the wild-type reporter construct induced in the absence of ncAAs.

**Fig. S29:**
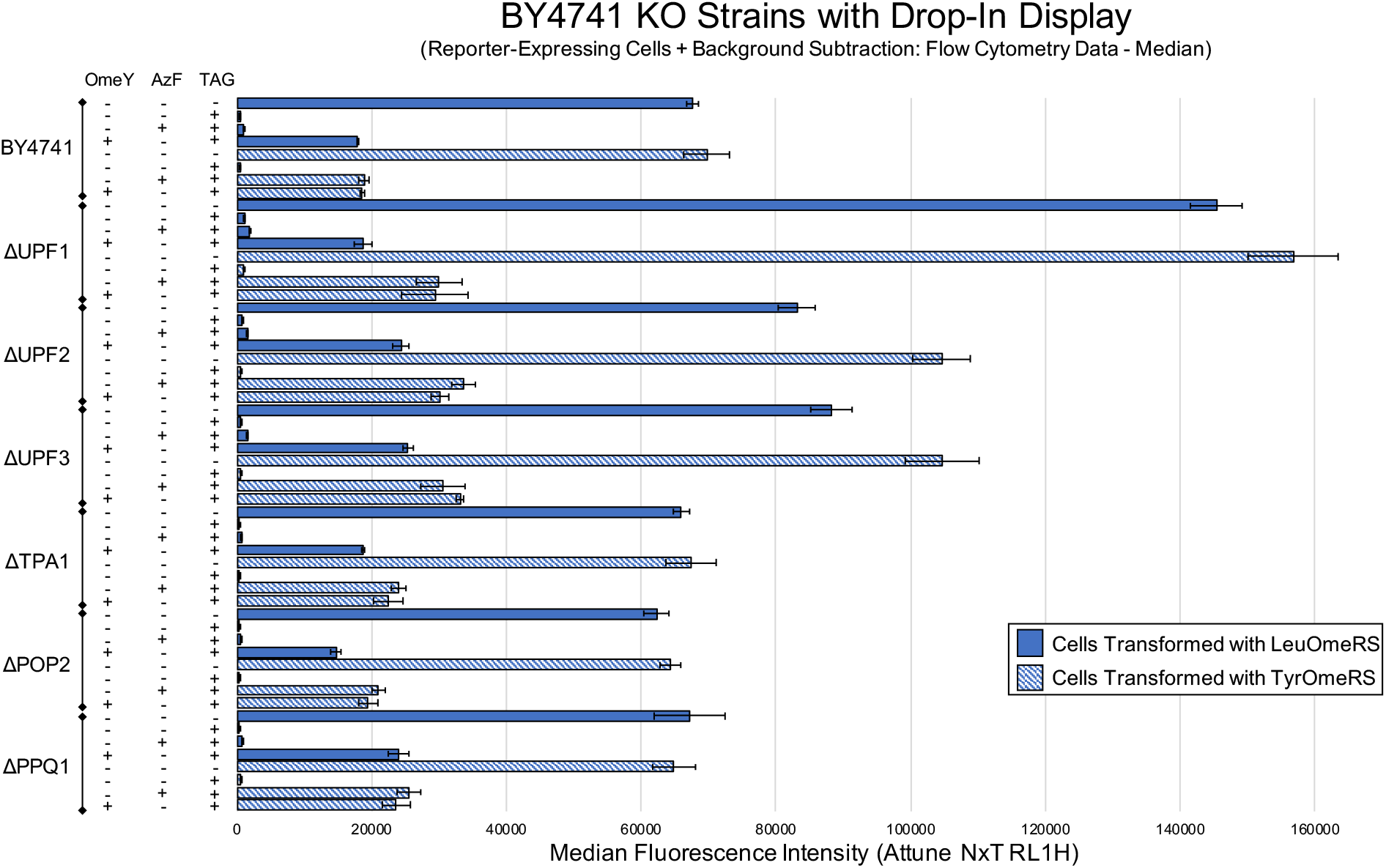
Alternate analysis of BY4741 and six single-gene knockout strains using the drop-in yeast display reporter. Measurements for the efficiency of ncAA incorporation calculated with median fluorescence intensity of reporter-expressing cells with background subtraction. The condition denoted by absence of ncAAs and a TAG codon is the wild-type reporter construct induced in the absence of ncAAs.

**Fig. S30:**
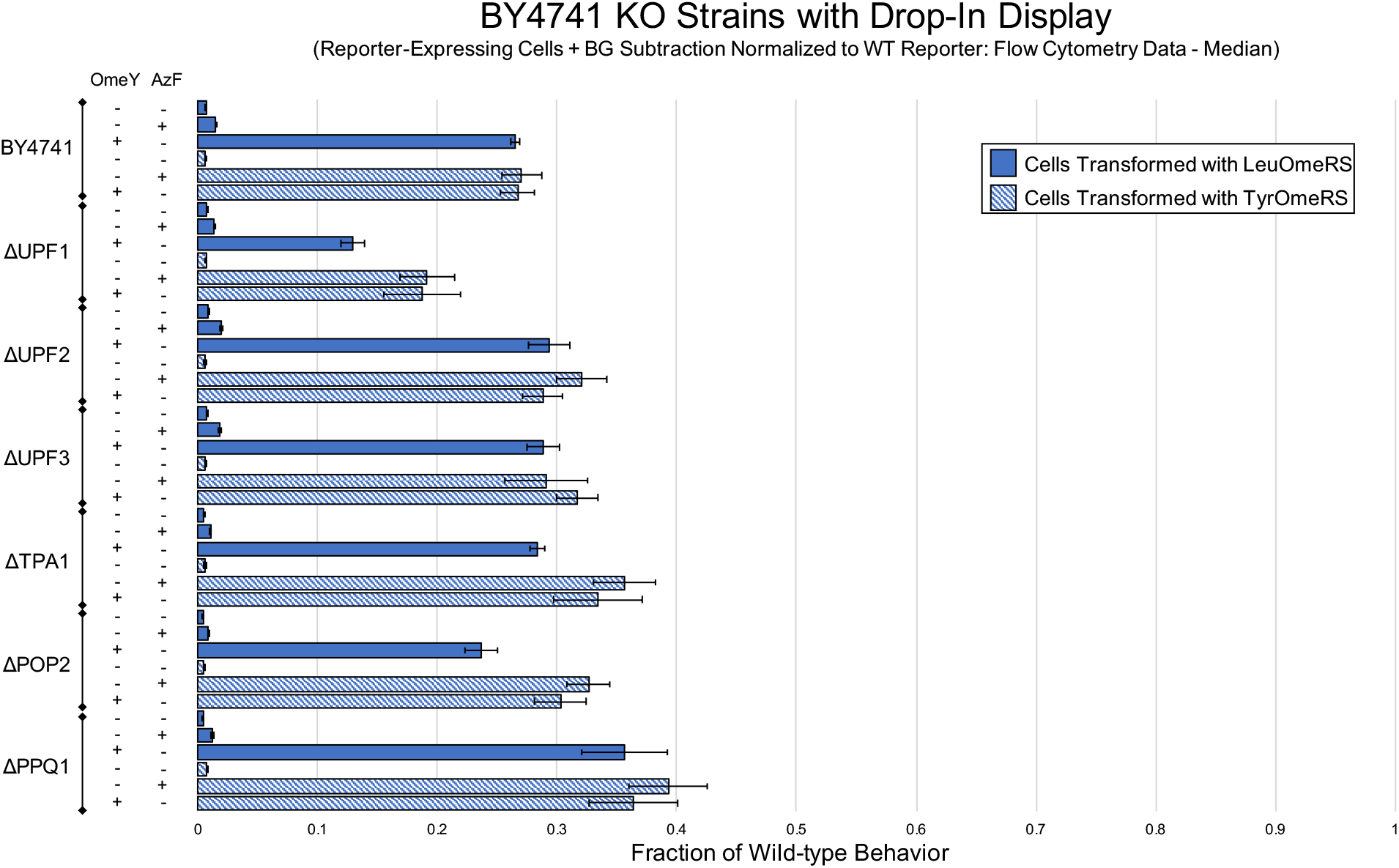
Alternate analysis of BY4741 and six single-gene knockout strains using the drop-in yeast display reporter. Measurements for the efficiency of ncAA incorporation calculated using the median fluorescence intensity of reporter-expressing cells with background subtraction and normalized to expression levels of cells expressing the wild-type reporter induced in the absence of ncAAs.

**Fig. S31:**
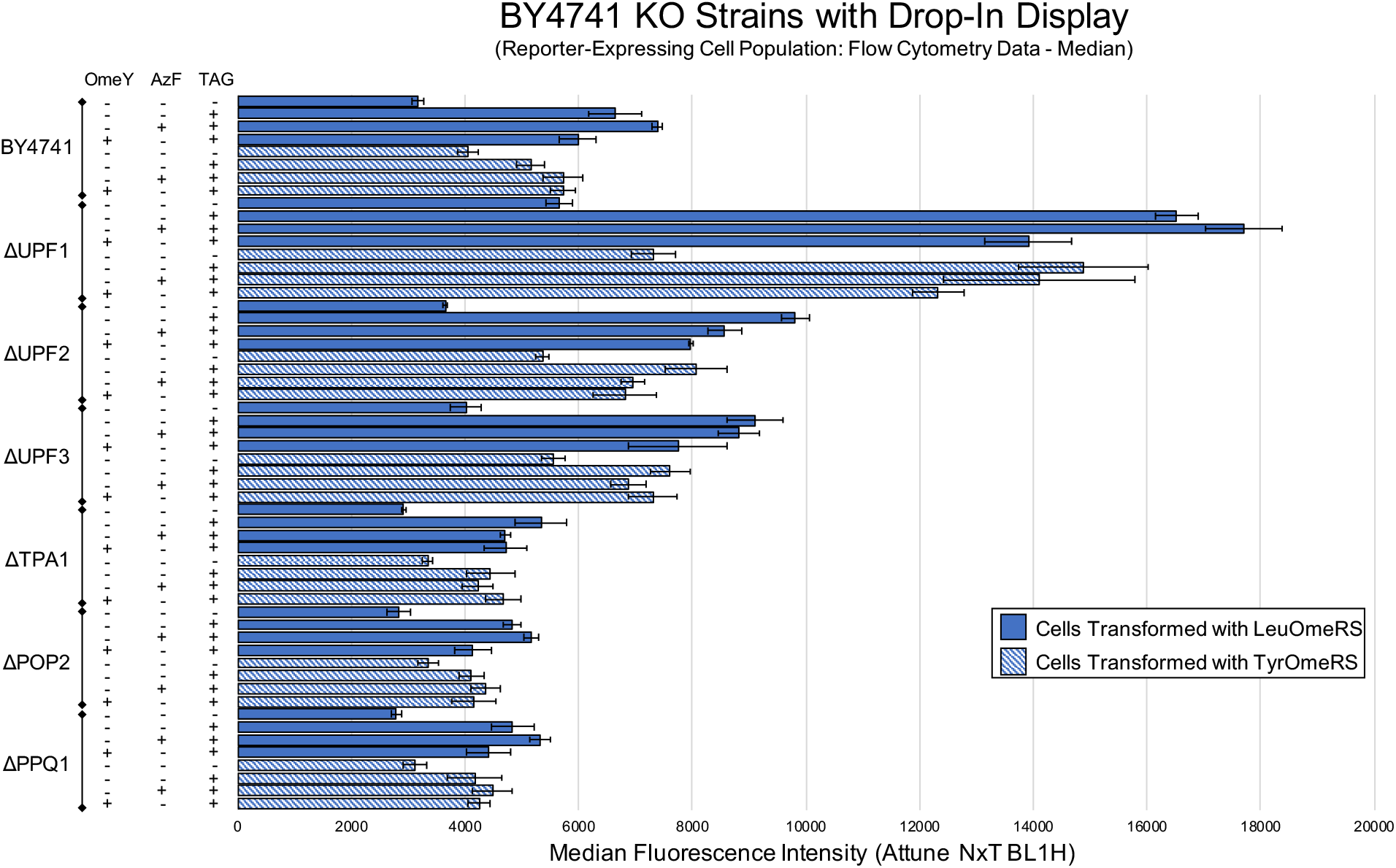
Measurement of reporter expression level in BY4741 and six single-gene knockout strains containing the drop-in yeast display reporter using the median fluorescence intensity of the cell population exhibiting above-background reporter expression. The condition denoted by absence of ncAAs and a TAG codon is the wild-type reporter construct induced in the absence of ncAAs.

**Fig. S32:**
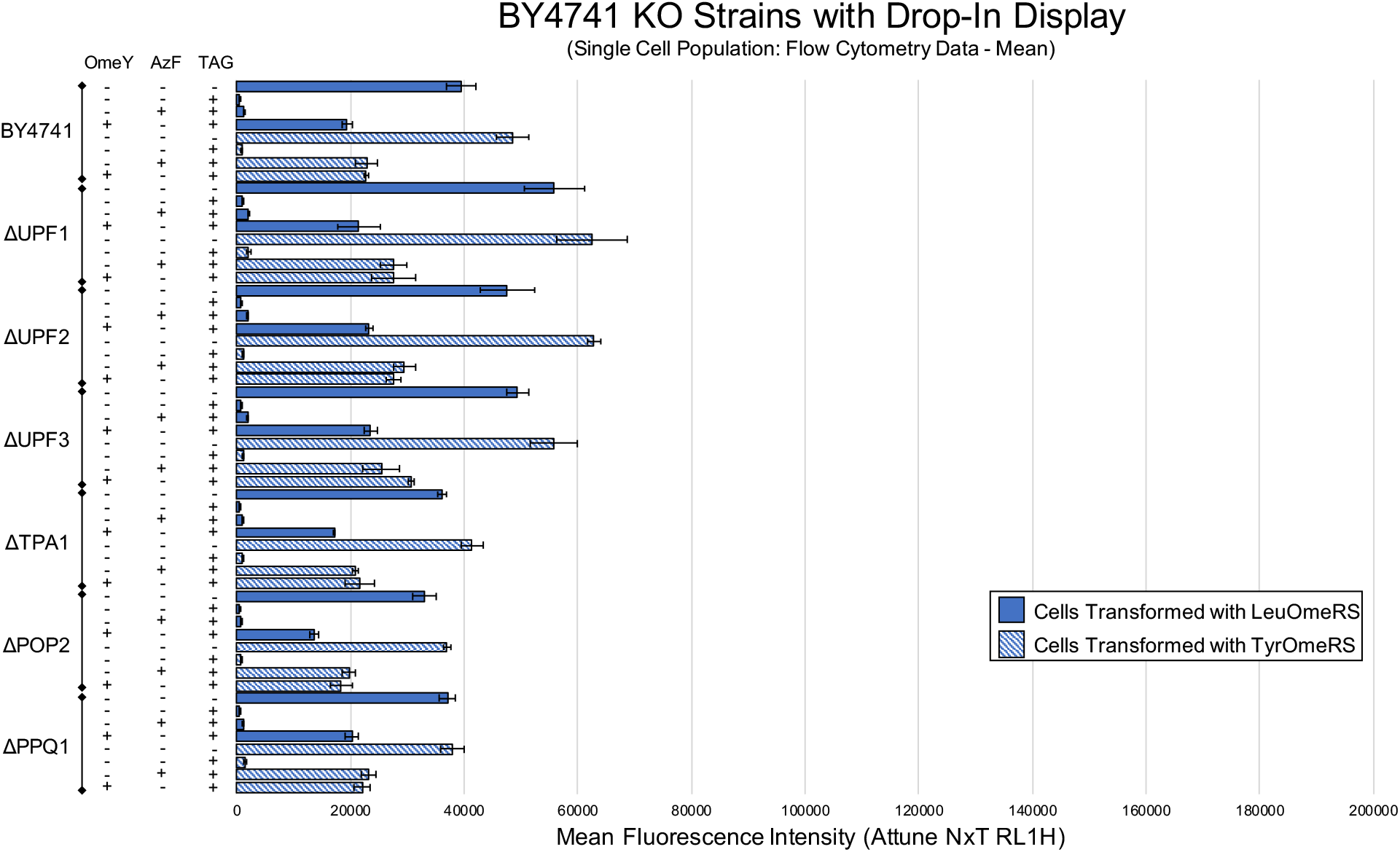
Alternate analysis of BY4741 and six single-gene knockout strains using the drop-in yeast display reporter. Measurements for the efficiency of ncAA incorporation calculated with mean fluorescence intensity using single cell population analysis. The condition denoted by absence of ncAAs and a TAG codon is the wild-type reporter construct induced in the absence of ncAAs.

**Fig. S33:**
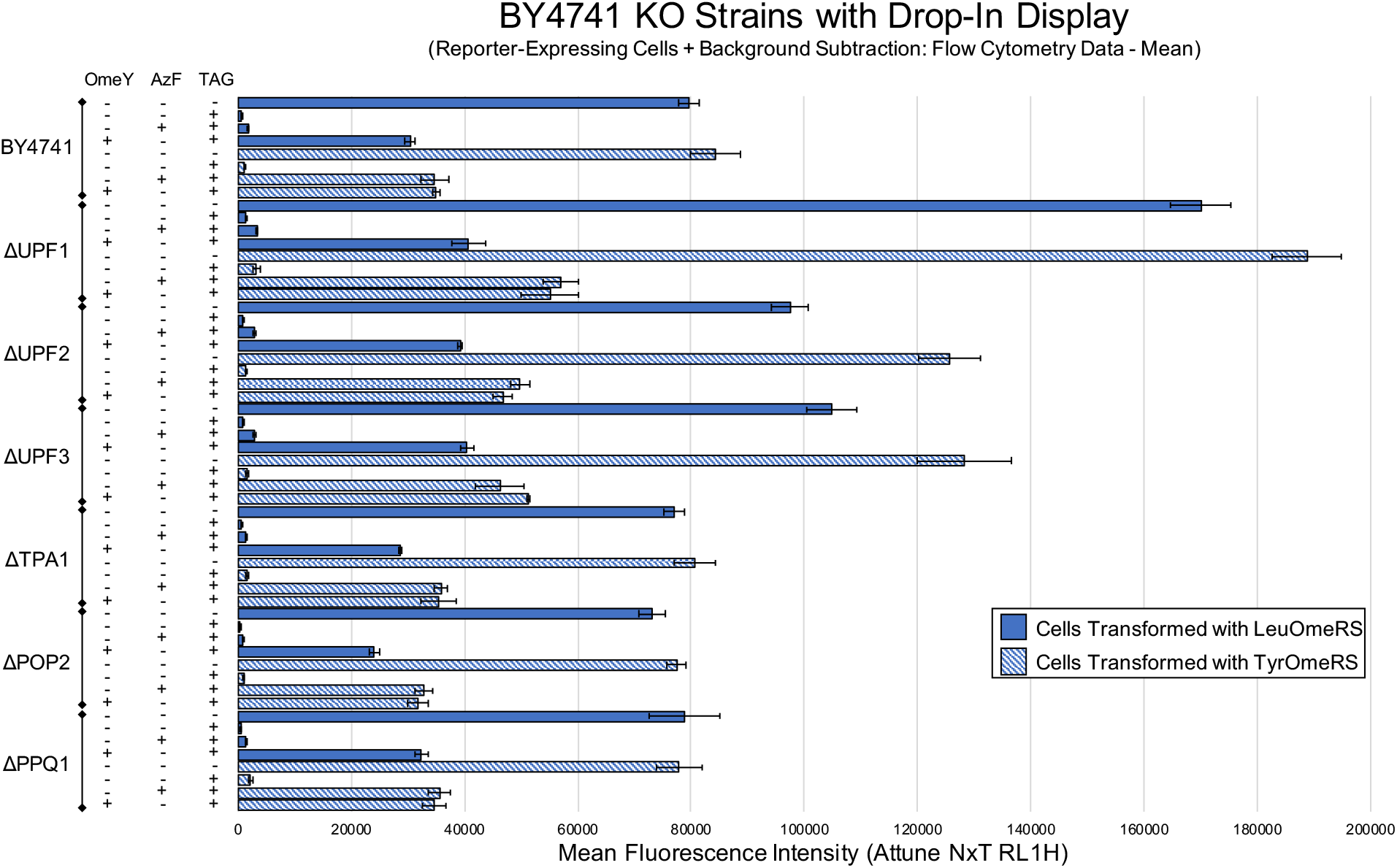
Alternate analysis of BY4741 and six single-gene knockout strains using the drop-in yeast display reporter. Measurements for the efficiency of ncAA incorporation calculated with mean fluorescence intensity using reporter-expressing cells with background subtraction. The condition denoted by absence of ncAAs and a TAG codon is the wild-type reporter construct induced in the absence of ncAAs.

**Fig. S34:**
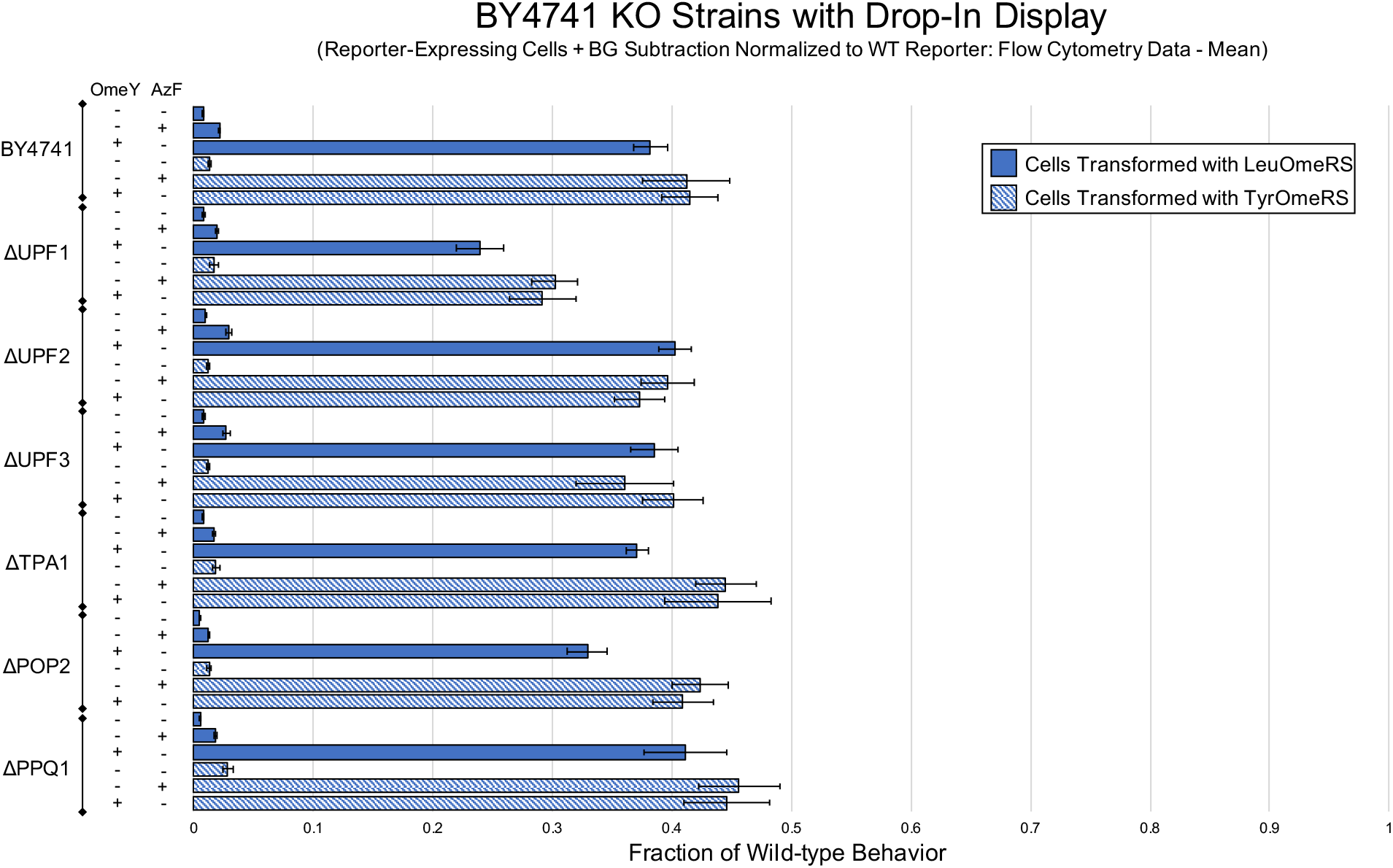
Alternate analysis of BY4741 and six single-gene knockout strains using the drop-in yeast display reporter. Measurements for the efficiency of ncAA incorporation calculated using the mean fluorescence intensity of reporter-expressing cells with background subtraction and normalized to expression levels of cells expressing the wild-type reporter in the absence of ncAAs.

